# Does more data solve the problem of deep support? A phylogenomic approach to clarifying evolutionary relationships in the *Lecanoraceae*

**DOI:** 10.64898/2025.12.18.694902

**Authors:** L.J. Li, C. Ivanovich-Hichins, S. LaGreca, L. Muggia, L. Myllys, S. Pérez-Ortega, G. Thor, L. Weber, E. Vakkari, C. Printzen, J. Gerasimova

**Author notes:** Corresponding authors: J. Gerasimova,; C. Printzen.

## Abstract

The family *Lecanoraceae* is one of the most diverse groups within *Lecanoromycetes*, yet its internal relationships have remained unresolved despite previous multilocus studies. Here we present the first phylogenomic reconstruction of the family, based on 1003 nuclear orthologous genes from 65 genome assemblies representing the major species groups and genera. Phylogenetic trees inferred from coalescent-based analyses, together with Quartet Sampling (QS) evaluation, confirm the monophyly of *Lecanoraceae* and recover several well-supported lineages. The *Lecanora subfusca* group is resolved as polyphyletic, comprising three major clades together with *Zeora* and *Vainionora*. Additional strongly supported lineages correspond to *Glaucomaria* s. lat., the MPRPS clade, and *Lecidella* s. lat., while *Bryonora* and *Frutidella* represent early-diverging members of the family. The *L. fuscescens* group and *Miriquidica* form a distinct lineage sister to *Lecanoraceae*, whereas *Ramboldia* (*Ramboldiaceae*) constitutes another sister family. Backbone support was generally improved, and QS analyses showed counter-support for only a few deep nodes, indicating a complex evolutionary history likely shaped by incomplete lineage sorting, hybridization, or rapid radiation. Our results demonstrate that genome-scale data improve the resolution of *Lecanoraceae*, but increasing the number of genes alone can not fully resolve the backbone relationships of the family.

## INTRODUCTION

*Lecanoraceae* Körb. (1855) is one of the largest families of the *Lecanoromycetes* (Kraichak *et al*. 2018), currently containing 35 genera and almost 3.800 published legitimate names, of which more than 700 species are currently accepted (Hyde *et al*. 2024, https://www.mycobank.org, accessed on 16 July 2025). Prior to Hafellner’s (1984) seminal work, circumscriptions of the family also included genera that are nowadays assigned to diverse classes and subclasses within Pezizomycotina outside Lecanoromycetidae, such as *Candelariella* Müll. Arg. (*Lichinomycetes*), *Phlyctis* (Wallr.) Flot, *Icmadophila* Trevis., and *Ochrolechia* A. Massal. (Ostropomycetidae). Although based solely on morphological evidence, with a strong focus on ascus apical structures, Hafellner’s concept of *Lecanoraceae* has, in general, stood the test of time. It included 17 names of genera (some of them only synonyms) of crustose to effigurate lichens with green-algal symbionts, apothecia with a thalline or rarely biatorine or lecideine margin, and *Lecanora*-type asci containing 8–16 simple and hyaline ascospores (Hafellner 1984).

Most of the species that are now assigned to *Lecanoraceae* traditionally belonged to the name-giving genus *Lecanora* Ach., nearly all containing algae in their apothecial margins, known as “lecanorine” apothecia. The focus on ascus apical structures subsequently led to the inclusion of genera with so-called “lecideine” margins, such as *Adelolecia* Hertel & Hafellner*, Carbonea* (Hertel) Hertel, *Lecidella* Körb., or *Miriquidica* Hertel & Rambold (Hertel 1983, Hafellner 1984, Hertel & Rambold 1987, 1995), thereby adding morphological diversity to an otherwise rather homogeneous family of lichenized *ascomycetes*. Molecular phylogenetic approaches have confirmed the wider circumscription of *Lecanoraceae* (Zhao *et al*. 2016), and in addition, have led to the recognition and segregation of several genera, some proposed earlier (e.g., Choisy 1929), and others created newly (e.g., Rodriguez-Flakus & Printzen 2014, Davydov *et al*. 2021). This separation of smaller, phenotypically diverging groups has meanwhile increased the number of genera in the family to 36 (Hyde *et al*. 2024).

However, the relationship between these genera and their circumscription within *Lecanoraceae* remained open. Based on DNA sequence data, several attempts have recently been made to reconstruct the phylogenetic relationships among some of these groups (Zhao *et al*. 2016, Medeiros *et al*. 2021, Svensson *et al*. 2022, Pérez-Ortega *et al*. 2023, Ivanovich *et al*. 2025b), typically facing trade-offs between taxon sampling and genetic loci coverage. Some studies employed a more comprehensive taxon sampling but included only a few genetic loci, generally resulting in weak phylogenetic support of the relationships between genera and species groups. For example, the two-locus phylogeny by Zhao *et al*. (2016) and the three-locus phylogeny by Pérez-Ortega *et al*. (2023) showed unsupported backbones with several well-supported clades at the generic level. On the other hand, the low taxon sampling in phylogenetic studies that used a higher number of gene loci presented a different challenge. For example, the phylogenies presented by Zhao *et al*. (2016), Medeiros *et al*. (2021), and Svensson *et al*. (2022), based on the same six gene loci, along with the findings of Ivanovich *et al*. (2025b) based on a partially different set of seven genes, showed considerably higher support for interior nodes—however, variation in taxon sampling resulted in sometimes conflicting results. For example, in Svensson *et al*. (2022), *Lecidella* formed a clade sister to the remaining *Lecanoraceae* (except for *Miriquidica*). In contrast, Ivanovich *et al*. (2025b) showed *Lecidella* as the closest lineage to the so-called MPRPS-clade (as defined by Medeiros *et al*. 2021), whereas Zhao *et al*. (2016) recovered it as closely related to the *Lecanora subfusca* and *Lecanora subcarnea* groups. The *Lecanora subfusca* group was inferred as monophyletic by Zhao et al. (2016), though this result may reflect limited taxon sampling. Medeiros *et al*. (2021) showed possible paraphyly of the *L. subfusca* group with the *L. subcarnea* group nested within it and relevant nodes lacking strong support. Ivanovich *et al*. (2025b) further supported this interpretation, recovering the *L. subfusca* group as distinctly non-monophyletic, forming a sister relationship to the resurrected genus *Zeora* (the former *L. symmicta* group). Additionally, *Bryonora* Poelt was strongly supported within the MPRPS-clade in Medeiros *et al*. (2021), whereas Ivanovich *et al*. (2025b) recovered its intermediate position among the *L. subfusca* group, *Zeora*, *Vainionora* and *Glaucomaria*. In Svensson *et al*. (2022), on the other hand, it formed a sister clade to *Lecanoraceae*, excluding the MPRPS clade.

Obtaining genome-scale data sets has recently become more feasible and has been used in phylogenetic studies on fungi, so far mostly at higher taxonomic levels (Shen *et al*. 2020, Li *et al*. 2021, Díaz-Escandon *et al*. 2022, Chen *et al*. 2023, He *et al*. 2024), employing several hundred to more than a thousand gene loci. It is not surprising that interior branches in phylogenetic trees are typically more robustly supported when they are based on a larger number of genes. In some cases, researchers discovered previously unrecognized lineages. For example, Díaz-Escandón *et al*. (2022) combined six morphologically diverse classes of *Ascomycetes* into *Lichinomycetes* based on almost 1300 single-copy orthologue genes. At lower taxonomic levels, genomic data sets have significantly enhanced our understanding of phylogenetic relationships within orders or families (Grewe *et al*. 2020, Kandemir *et al*. 2022, Liimatainen *et al*. 2022, Qu *et al*. 2025). However, an increase in data, including more genetic loci, does not always lead to unambiguous and better-supported phylogenies. For example, Qu *et al*. (2025) found evidence for reticulate evolution caused by hybridization and incomplete lineage sorting. Additionally, Grewe *et al*. (2020) observed that including a higher number of fast-evolving genes in their data matrix for *Parmeliaceae* resulted in low support values. In this study, we investigate whether metagenomic data can help us to infer a well-supported phylogenetic backbone for the family *Lecanoraceae*. Our goal was to include as many key representatives of the species groups and genera recently segregated from *Lecanora* or assigned to the family *Lecanoraceae* as possible. Since most of the species in *Lecanoraceae* are small and difficult to cultivate in the laboratory, we used the whole genome amplification (WGA) approach as a simple and efficient method for generating DNA samples with sufficiently high concentration and quality for short-read next-generation sequencing, as described by Ivanovich *et al*. (preprint).

## MATERIALS AND METHODS

### Sample selection and phenotypic study

We selected 58 species from the family *Lecanoraceae*, including representatives from 22 species groups within *Lecanora* and related genera. These include: *Adelolecia* Hertel & Hafellner, *Bryonora* Poelt, *Frutidella* Kalb, *Glaucomaria* M. Choisy (*L. carpinea*-*rupicola* group), *Lecanora* s. str. (*L. subfusca* group), the *L. albella* group, the *L. fuscescens* group, the *L. intumescens* group, *Lecanoropsis* M. Choisy ex Ivanovich (the former *L. saligna* group), “*Lecidea*” *tuberculifera, Lecidella*, *Miriquidica, Myriolecis* Clem. (the former *L. dispersa* group), *Nimisora* Pérez-Ort., M. Svenss. & J.C. Zamora, *Protoparmeliopsis* M. Choisy (the former *L. muralis* group), *Ramboldia* Kantvilas & Elix*, Rhizoplaca* Zopf, *Straminella* M. Choisy (the former *L. varia* group), *Vainionora* Kalb, *Verseghya* S.Y. Kondr., Lőkös & Hur, *Zeora* Fr. (the former *L. symmicta* group), and *Xanthosyne* Lendemer, R.C. Harris, Brodo & McMullin.

Our sampling includes the type species of the genera *Adelolecia*, *Frutidella*, *Glaucomaria*, *Lecanora*, *Miriquidica, Nimisora*, *Protoparmeliopsis*, *Straminella*, and *Xanthosyne*. The generic type of *Verseghya*, *V. klarae* S. Y. Kondr. *et al*., is probably a synonym of *V. thysanophora* included in this study (see tree in Kondratyuk *et al*. 2019). The representatives of *Lecanoropsis*, *Myriolecis*, *Ramboldia*, *Rhizoplaca*, and *Zeora* included in our data set are phylogenetically closely related to the type species of these genera (Śliwa *et al*. 2023, Ivanovich *et al*. 2025a, 2025b). No fresh material was available for the type species of *Carbonea*, *C. atronivea* (Arnold) Hertel, and *Lecidella*, *L. viridans* (Flot.) Körb. and these species have never been included in any phylogenetic study. Nevertheless, *C. vorticosa* (Flörke) Hertel and *L. elaeochroma* (Ach.) M. Choisy included here are morphologically similar enough to the type species to assume that in the context of a family-wide phylogeny, their position will not be far from the hypothetical one of the generic types. A complete list of selected taxa is presented in Table S1, which includes Herbarium Number, Locality, and GenBank accession number. We requested nine specimens of *Lecanora* (NY) from corresponding genome assemblies from GenBank for morphological examination.

Macrophotographs of representatives of the included groups were taken using a Zeiss Axio Zoom V16 microscope (Zeiss, Jena, Germany). External morphological characters were studied on air-dried material using a stereomicroscope (Zeiss Stemi SV11, Zeiss, Jena, Germany). Anatomical features were investigated using a light microscope Zeiss Axioskop 2 plus (Zeiss, Jena, Germany) on transverse sections of apothecia and thalli, cut with a freezing microtome Zeiss HYRAX KS 34 (Zeiss, Jena, Germany) to 16–20 µm thickness and mounted in water, Lugol’s iodine solution (I), and lactophenol cotton blue (LCB). The specimens examined are deposited in the Herbarium H, FR, NY, O, HMAS, KUN, UPS, and the corresponding voucher information is available in Table S1.

### Dataset construction

The dataset consisted of a total of 65 genome assemblies from the family *Lecanoraceae*: 48 assemblies were newly generated for this study, two were provided by co-authors, eleven *Lecanora* assemblies were downloaded from GenBank (Table S1), and four GenBank assemblies from sister families were used as an outgroup: *Gypsoplaca macrophylla* (Zahlbr.) Timdal, *Letharia vulpina* (L.) Hue, *Parmelia squarrosa* Hale, and *Platismatia glauca* (L.) W.L. Culb. & C.F. Culb. We selected a high-quality chromosome assembly of *Platismatia glauca* (GCA_963556305.1) as the reference genome for our dataset. All assembly details are listed in Table S1. We kept the original taxon names from NCBI, regardless of their current taxonomic status, unless the specimens were examined microscopically by us.

*Japewia tornoensis* (Nyl.) Tønsberg, *Lecanora cancriformoides* Zahlbr., *L. cinereofusca* H. Magn., *L. subcarnea* (Lilj.) Ach., *Palicella glaucopa* (Hooker f. & Taylor) Rodr. Flakus & Printzen, *Sedelnikovaea baicalensis* (Zahlbr.) S.Y. Kondr., M.H. Jeong & Hur and an unidentified species of *Vainionora* were initially included in our sampling dataset, but due to insufficient sequencing depth, likely caused by the low amount of DNA retained, only a few non-orthologous genes remained. In addition, a specimen included to represent the *L. polytropa* group, turned out to be *Myriolecis* cf. *dispersa* due to a laboratory error. No fresh material was available for the other 16 out of the 36 genera within the *Lecanoraceae* (Hyde *et al*. 2024). Thus, the following genera and groups were not represented in our dataset: *Ameliella* Fryday & Coppins, *Bryodina* Hafellner, *Carbonicola* Bendiksby & Timdal, *Cladidium* Hafellner, *Claurouxia* D. Hawksw., *Clauzadeana* Cl. Roux *, Edrudia* W.P. Jordan, *Japewia* Tønsberg, *Japewiella* Printzen, the *L. polytropa* group, *Maronina* Hafellner & R.W. Rogers, *Maronora* Kalb & Aptroot, *Omphalodina* M. Choisy, *Palicella*, *Psorinia* Gotth. Schneid., *Pulvinora* Davydov, Yakovch. & Printzen, *Punctonora* Aptroot, *Pyrrhospora* Körb., *Sagema* Poelt & Grube, and *Traponora* Aptroot.

### Sample preparation for the whole-genome amplification

Specimens were cleaned with a brush to minimize contamination. Horizontal cross-sections of the apothecia were created using a HYRAX KS 34 cryotome (Zeiss, Jena, Germany), following the protocol described by Ivanovich *et al*. (preprint). In detail, the first several slices, approximately 20 μm thick each, from the upper part of the hymenium were discarded until the visible algal layer was reduced, thereby eliminating the algal layer and reducing the risk of contamination from non-target fungi present on the apothecia’s surface. Additionally, all tools and the cryotome knife were carefully disinfected with bleach before and after each step to prevent cross-contamination. The subsequent three cross-sections of the apothecia, approximately 10 μm thick, were placed directly into 1.5 ml Eppendorf tubes and stored at -20 °C for DNA amplification. DNA extractions of *Lecanora cadubriae* and *L. circumborealis,* provided by our collaborators in Helsinki, were performed using samples from several apothecia (at least 1 mg) that were freshly collected. The apothecia were stored in 1.5 ml microcentrifuge tubes at +4 °C for 46 hours before extraction.

### DNA amplification, isolation, and sequencing

#### Illumina NovaSeq X Plus Series (PE150)

High-molecular-weight genomic DNA was amplified using the Qiagen Repli-G Single-Cell Kit (Qiagen GmbH, Düsseldorf) according to the manufacturer’s protocol with one modification: the incubation time was extended to 16 hours at 30°C overnight to ensure sufficient DNA concentration for library preparation. DNA concentrations and quality were assessed using the TapeStation 2200 System (Agilent Technologies, Santa Clara, CA, USA). A total concentration of 80 µg (between 0.1 and 0.2 μL in volume) was used for 150 bp paired-end libraries using the Illumina DNA Prep kit and sequenced on the NovaSeq X Plus Series, generating 10 Gb of raw data per sample, following the Plant and Animal Whole Genome Library Preparation (350 bp) protocol performed by Novogene GmbH (Planegg-Martinsried, Munich, Germany).

#### PacBio data

The samples were flash-frozen in liquid nitrogen and coarsely ground on dry ice using a micropestle. This step was followed by manual homogenization in an extraction buffer composed of 50 mM Tris-HCl (pH 7.2), 50 mM EDTA (pH 8.0), 3% (w/v) SDS, and 1% (v/v) 2-Mercaptoethanol at room temperature. After homogenization, 0.2 mm diameter glass beads were added, and the samples were vortexed at 3200 rpm for 2 minutes.

Cell lysis was performed at 65 °C for one hour, with intermittent mixing at 750 rpm for 40 seconds every 10 minutes. RNase A treatment (Thermo Scientific EN0531) was then conducted at a working concentration of 600 μg/mL at 37 °C for 30 minutes, using the same intermittent mixing protocol. Total DNA was purified from the lysate through phenol/chloroform extraction followed by isopropanol precipitation. High-molecular-weight DNA was enriched using the MagAttract HMW DNA kit (Qiagen 67563), according to the manufacturer’s protocol for tissue samples, but with a modified lysis time of 1 hour. DNA purity and fragment size were measured with a NanoDrop 2000 spectrophotometer (Thermo Fisher Scientific Inc., Waltham, MA, USA) and an Agilent 2200 TapeStation System (Agilent Technologies, Santa Clara, CA, USA), respectively. DNA concentration was determined using a Qubit 4 fluorometer with the dsDNA HS assay kit (Invitrogen Q32854).

PacBio HiFi SMRTbell sequencing libraries were prepared according to the manufacturer’s ultra-low DNA input protocol and sequenced on the Sequel II platform at the DNA Sequencing and Genomics Laboratory, Institute of Biotechnology, University of Helsinki. HiFi reads were generated using SMRTlink v11.1. The assemblies were performed by the DNA Sequencing and Genomics Laboratory at the Institute of Biotechnology, University of Helsinki, using PacBio’s Genome Assembly protocol. The default parameters from the same protocol were utilized to generate and demultiplex HiFi reads and to run the Genome Assemblies application (see p. 56 of PacBio’s Genome Assembly protocol, Supplementary Material S6).

### Quality assessment

The primary quality control checks on raw sequence data, performed with FastQC v0.12.1 (Andrews 2010), showed that all reads had a quality above 20. The duplicated reads were removed using ParDRe v2.1.5 (González-Domínguez & Schmidt 2016). The following steps for cleaning adapters and low-quality reads were performed using Captus v1.0.1 (Ortiz *et al*. 2023; https://edgardomortiz.github.io/captus.docs/index.html), with the average PHRED quality score set to its default value. The quality of the deduplicated and cleaned reads was assessed with FastQC prior to assembly and is presented in the Supplementary material (S1-S2).

### Genome assembly

The assembly step was carried out using Captus v1.0.1 (Ortiz *et al*. 2023). Based on the cleaned reads obtained from the previous step, Captus performs *de novo* assembly utilizing MEGAHIT. An HTML report summarizing the assembly statistics generated during this step can be found in the Supplementary material (S3). To exclude low-coverage contigs, the minimum contig depth was set to 5 (--min_count 5), and the minimum contig length was set at 500 base pairs (--min_contig_len 500).

To extract the genes of interest from our metagenome assemblies, we provided the coding sequences of the reference assembly (captus extract -a 02_assemblies -f *.fasta -n Platismatia_glauca.cds-transcripts.fasta --nuc_min_identity 70 --nuc_min_coverage 50 --nuc_min_score 0.13 --ram 100 --threads 20 --concurrent 2 -o 03_extractions). We run a predict step using funannotate v1.8.15 (Palmer *et al*. 2020) on a chromosome-high-quality assembly of *Platismatia glauca* (GCA_963556305.1) to obtain a cds-transcripts file.

The minimum percentage of identity (--nuc_min_identity) required for a match with the reference protein was set to 70% for *Lecanoraceae* and closely related families. A stricter threshold of 80% was applied to the outgroup, as it belongs to the same family or a closely related one. The minimum percentage of coverage of the reference protein required for a hit to be retained was set to 50 (--nuc_min_coverage 50), with a minimal score set to the default of 0.13 (default), meaning that at least 13% of the protein query must be found on one contig.

An HTML report summarizing the extract statistics can be found in the Supplementary material (S4).

### Alignment, trimming, and phylogeny reconstruction

The extracted markers were aligned with MAFFT as a part of the Captus pipeline (captus align -e 03_extractions_all -m NUC -f NT,GE --max_paralogs 5 --align_method mafft_auto --clipkit_method smart-gap --filter_method none --ram 200 --threads 20 --concurrent 2) with a minimum number of samples in a marker to proceed (--min_samples) set to 25. Captus initiates this step by gathering all the markers across the extracted samples and building a FASTA file for each marker, resulting in 4649 homologous gene alignments. We used all the recovered sequences, including paralogs, for inferring a species tree using a coalescent-based approach that accounts for paralogy. To reduce uncertainty in tree estimation due to insufficient data and reduce the impact of missing data, we collapsed the branches with low support with a minimum of 70% threshold (Morales *et al*. 2022; https://bitbucket.org/dfmoralesb/target_enrichment_orthology/src/master/scripts/collapse_branches_bs_multiphylo.py).

We estimated a maximum likelihood (ML) tree for each alignment using IQ-TREE v2.2.6 (Minh *et al*., 2020), searched for the best model using ModelFinder, which is implemented within IQ-TREE (Kalyaanamoorthy *et al*. 2017), and performed 1000 ultrafast bootstrap replicates to assess clade support. The output tree files from the IQ-TREE runs were combined into a single file. We then inferred a multigene coalescent species tree using the resulting gene tree with ASTRAL-PRO 3 v1.20.3.6 (Zhang *et al*. 2020). The resulting phylogeny was inferred to retain all samples, even when only a limited number of genes were available for some of them, and allowed us to evaluate the preliminary results—the resulting phylogeny is presented in the Supplementary material (S5).

### Orthology inference of nuclear loci

Orthology inference was performed following a modified version of the methods described in Morales-Briones *et al*. (2022, https://bitbucket.org/dfmoralesb/target_enrichment_orthology), which used the tree-based ‘monophyletic outgroup’ (MO) approach described in Yang and Smith (2014). The MO method searches for clusters with monophyletic ingroups rooted at the outgroups in the homologue trees, discarding those with duplicated taxa in the outgroups. Subsequently, it infers the orthologues from root to tip, keeping the orthologue subtree with the most taxa. First, the output 4649 homologous trees were masked for both mono- and paraphyletic tips that belonged to the same taxon, keeping the tip with the most unambiguous characters in the trimmed alignment, using the modified *mask_tips_by_taxonID_transcripts.py* script. Then, we trimmed spurious tips using the *tree_shrink_1.3.9_wrapper.py* script. We filtered homologs with monophyletic, non-repeating outgroups and rerooted and cut paralogs from the root to the tip using *prune_paralogs_MO_1to1_MO.py*. We visualized the final occupancy stats to check if any taxa have an unusually low number of genes in the orthologs and decided on the minimum number of taxa for the supermatrix. For more details on each step of the pipeline, please refer to https://bitbucket.org/dfmoralesb/target_enrichment_orthology/src/master/. The extracted markers were aligned using MAFFT v7.525 --auto (Katoh *et al*. 2019) and trimmed with Clipkit v2.3 --smart-gap (Steenwyk *et al*. 2020). Once we had clean alignments, we chose the minimal cleaned alignment length, set to 1000, and the minimal number of taxa, set to 48 of each ortholog, to include in the supermatrix (*concatenate_matrices_phyx.py*). The resulting dataset comprised 1003 orthologous gene alignments.

### Phylogenetic reconstruction

We used coalescent-based methods to reconstruct the phylogeny of *Lecanoraceae*. First, we estimated a ML tree for all 1003 nuclear loci produced using the clean orthologue alignments with IQ-TREE v.2.2.6 (Minh *et al*. 2020). We searched for the best model using ModelFinder, which is implemented within IQ-TREE (Kalyaanamoorthy *et al*. 2017), and performed 1000 ultrafast bootstrap replicates to assess clade support.

The output tree files from the IQ-TREE runs were combined into a single file. We inferred the species tree using the 1003 individual ML orthologue trees with default ASTRAL parameters and assessed branch support with local posterior probability (LPP, Sayyari and Mirarab 2016).

### Gene trees discordance estimation

We followed the procedure outlined by Lagou *et al*. (2024) to evaluate conflict among nuclear gene trees at each node of the inferred species tree. Specifically, we estimated the number of conflicting and concordant bipartitions using Phyparts (Smith *et al*. 2015). For this analysis, we utilized individual maximum likelihood (ML) orthologue trees and established a threshold of at least 70% BS support to classify a node as informative. The results from Phyparts were visualized with a phypartspiecharts_missing_uninformative.py script to account for missing data and uninformative nodes (https://bitbucket.org/dfmoralesb/target_enrichment_orthology) in Python v.3.10.10. This script generated piecharts, considering instances where the input trees differed in the number of tips (i.e., missing data).

Additionally, we applied Quartet Sampling (QS, Pease *et al*. 2018) to differentiate between nodes that lacked support and those that exhibited conflict in the species tree. For a given phylogeny, each observed internal tree branch partitions the tree into four nonoverlapping subsets of taxa, which can exist in three possible relationships: the concordant relationship that matches the configuration in the given topology, and two alternative discordant configurations. The QS method repeatedly and randomly samples one taxon from each of the four subsets and then evaluates the likelihood all three possible phylogenies given the sequence data (either the full alignment or a randomly sampled gene partition) for the randomly selected quartet spanning that particular branch (Pease *et al*. 2018). QS estimates branch support and conflict by sampling quartets from the species tree and the corresponding concatenated alignment. QS calculates the frequency of the three possible topologies for each node, enabling a simultaneous evaluation of three factors: the consistency of information (Quartet Concordance, QC), the presence of secondary evolutionary history (Quartet Differential, QD), and informativeness of internal nodes (Quartet Informativeness, QI). Each internal branch was annotated with a set of three scores (Quartet Concordance/Quartet Differential/Quartet Informativeness: QC/QD/QI), providing complementary information to the BS and LPP of the inferred species tree. A detailed explanation of the QS methodology and interpretation of the result values can be found in the original paper by Pease *et al*. (2018). We ran 10000 QS replicates with RAxML-NG (Kozlov *et al*. 2019) as the ML inference tool. The results were plotted using R (R Core Team 2023), with the values of QC on each node colour-coded and annotated with the rest of the estimated values (https://bitbucket.org/yanglab/conflict-analysis/src/master/). The final phylogeny was visualized using FigTree v1.4.4 and a custom R script, with additional annotations added in Adobe Illustrator v29.5.1.

## RESULTS

### Phylogenetic relationships

Our data set constitutes the most comprehensive gene and taxon sampling for genus-level groups within the family *Lecanoraceae* to date, comprising approximately half of the genera and species groups that have so far been assigned to this family. The *Lecanoraceae* clade comprises the following subclades (from top to bottom in Fig. 1): *Lecanora subfusca* group 2, *L. subfusca* group 3, *Zeora*, and *L. subfusca* group 1; *Vainionora* (represented by *Lecanora markjohnstonii* And. Stewart, E. Tripp & Lendemer, see discussion); a clade comprising *Glaucomaria* (the former *L. rupicola* group), *L. intumescens, Verseghya* and two specimens of *L. albella* (“*Glaucomaria* s. lat.”); the MPRPS clade comprising *Myriolecis* (the former *L. dispersa* group), *Protoparmeliopsis*, *Rhizoplaca*, *Carbonea*, *Straminella* and *Lecanoropsis*; a clade comprising *Xanthosyne*, *Lecidella*, and *Nimisora* (“*Lecidella* s. lat.”); and a clade containing *Bryonora* and *Frutidella*. All these clades can be interpreted as the core of *Lecanoraceae*. Sister to the core *Lecanoraceae* are two nodes. One of these includes *L. cadubriae* (representing the *L. fuscescens* group) and *Miriquidica,* the other one includes *Ramboldia*. These two groups are provisionally called *Miriquidicaceae* and *Ramboldiaceae* in this study.

**Figure 1.**
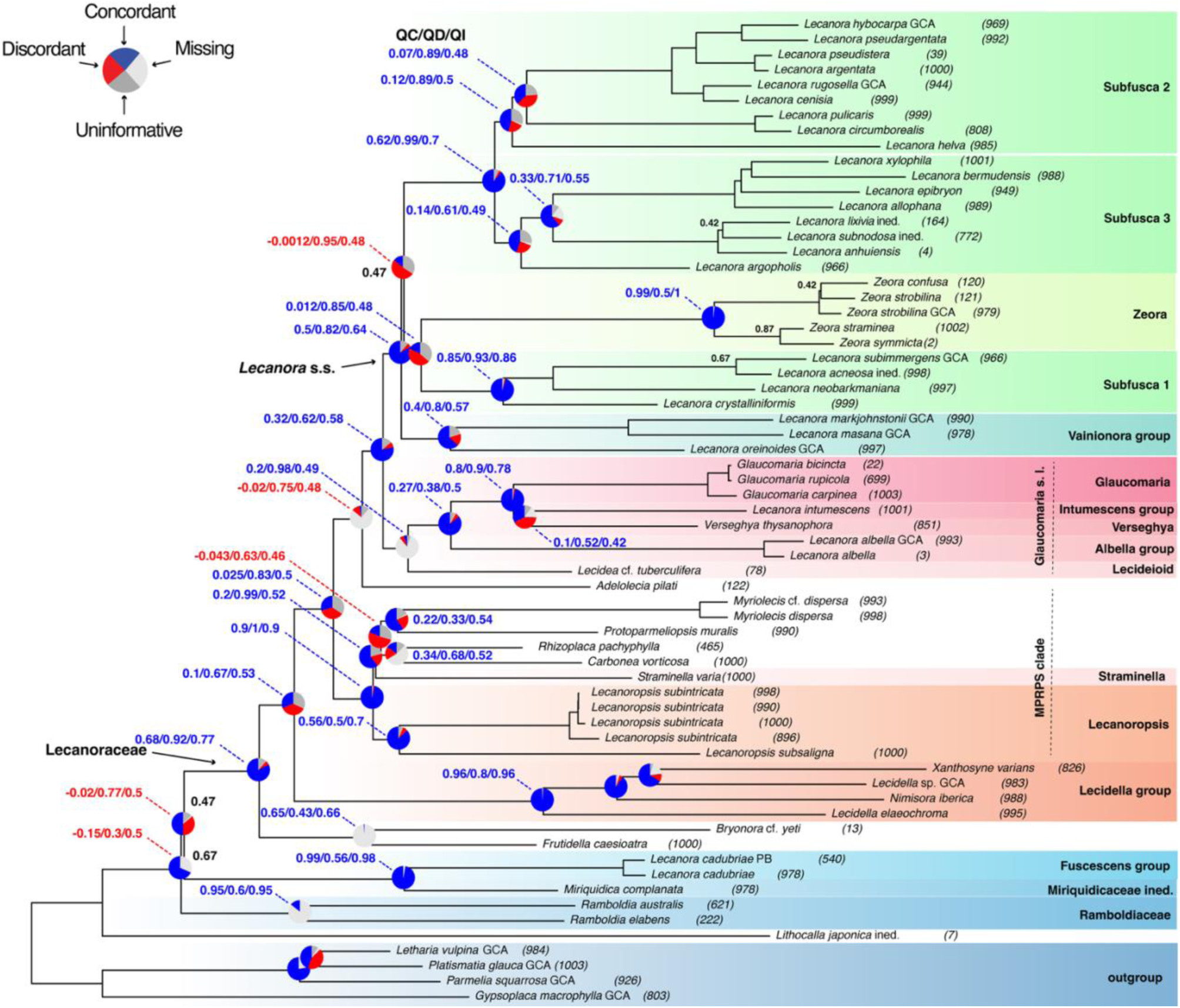
The coalescence-based phylogeny of the family *Lecanoraceae*, inferred from 1003 orthologous genes using ASTRAL. Local posterior probabilities are shown above the branches when < 0.95. The number of orthologs per species given in the parentheses. Colours in the pie charts: blue denotes the proportion of concordant gene tree topologies, dark grey denotes the proportion of gene trees with the main alternative topology, red denotes the proportion of gene trees with the remaining discordant topologies, light grey denotes the proportion of gene trees with missing taxa and dark grey denotes the proportion of uninformative gene trees. The outgroup consists of *Gypsoplaca macrophylla*, *Letharia vulpina*, *Parmelia squarrosa*, and *Platismatia glauca*.

Several taxa with uncertain taxonomic placement are dispersed across the phylogenetic tree. *Lecanora masana*, which is morphologically similar to the *L. subfusca* groups, is sister to *Vainionora*. The phenotypically distinct *L. oreinoides* forms a clade with both *L. masana* and *Vainionora*. The species referred to as “*Lecidea*” *tuberculifera* is sister to *Glaucomaria* s. lat. *Adelolecia pilati* (Hepp) Hertel & Hafellner is sister to the clade that combines the *L. subfusca* groups, *Zeora*, *Vainionora* and *Glaucomaria* s. lat. Notably, a specimen originally identified as a member of the *L. polytropa* group is, in fact, an undescribed species (*Lithocalla japonica* ined.) forming a sister clade to the families *Lecanoraceae*, *Miriquidicaceae* and *Ramboldiaceae*. The lecideoid species of *Carbonea*, *Adelolecia,* “*Lecidea*”*, Xanthosyne, Lecidella, Nimisora*, and *Frutidella* are polyphyletic belonging to separate clades throughout the tree.

### Statistical support

High LPP and QS supports were recovered for most internal and external branches in the inferred phylogenies, except for some branches along the backbone, discussed below. The monophyly of *Lecanoraceae* is highly supported (LPP = 100). Phyparts identified 886 informative concordant genes out of a total of 1003, and the quartet sampling analysis yielded high support values (QC/QD/QI: 0.68/0.92/0.77), based on a substantial proportion of quartet samples (7076). Maximum support, high concordance and congruence between the phylogenies were also inferred for the close relationship between the *L. subfusca* clades 2 and 3 (LPP = 100, Phyparts: 925/1003, QC/QD/QI: 0.62/0.99/0.7) and the one between *Zeora* and *L. subfusca* clade 1 (LPP = 100, Phyparts: 270/1003, QC/QD/QI: 0.012/0.85/0.48). While only 270 out of 1003 gene trees were concordant regarding this last node the remaining 732 gene trees did not display any consistent topology. This placement aligns with the classification of Ivanovich *et al*. (2025a, 2025b), which suggests that *Zeora* is a sister clade to *L. subfusca* clade 1 based on Sanger sequences. *Vainionora*, represented in our phylogeny by *L. markjohnstonii* (see discussion), was recovered as sister to *L. masana* and *L. oreinoides* with high support (LPP = 100, Phyparts: 712/1003, QC/QD/QI: 0.4/0.8/0.57). The core group of *Lecanora* comprises the three *L. subfusca* clades, *Zeora* and *Vainionora*, together forming a highly supported clade with high support (LPP = 100, Phyparts: 884/1003, QC/QD/QI: 0.5/0.82/0.64).

The sister relationship of this clade with the clade comprising *Glaucomaria* s. lat. and “*Lecidea*” *tuberculifera* is also highly supported (LPP = 100, Phyparts: 840/1003, QC/QD/QI: 0.32/0.62/0.58), as are the MPRPS clade (LPP = 100, Phyparts: 390/1003, QC/QD/QI: 0.25/0.83/0.5), and *Lecidella* s. lat. (LPP = 100, Phyparts: 314/1003, QC/QD/QI: 0.1/0.67/0.53). Two clades, *Bryonora* and *Frutidella,* formed sister lineages to the rest of *Lecanoraceae* with high support (LPP = 100, Phyparts: 886/1003, QC/QD/QI: 0.68/0.92/0.77) supporting the classification of Poelt (1983) and Kalb (1994), who included these two genera in *Lecanoraceae*.

### Discordance

Several nodes in the ML tree received counter-support in the quartet sampling analysis. One of these clades comprises *Lecanora* s. str., including *L. subfusca* clade 1, *L. subfusca* clade 2, *L. subfusca* clade 3, and *Zeora* (BS=47, Phyparts: 145/1003, QC/QD/QI: -0,0012/0.95/0.48). In this case, the countersupport derives from uncertainty about the position of *Vainionora* in relation to these groups. The relevant node in Fig. 1 is supported by 1540 quartet samples, while 1733 supported an alternative topology, in which *Vainionora* formed a sister clade to *Zeora* and *L. subfusca* clade 1 (Fig. 2A). Although the phylogenetic analyses revealed a high degree of discordance for *Zeora* as a sister clade of *L. subfusca* group 1 (QC/QD/QI: 0.012/0.85/0.48, with 180 informative concordant genes out of a total of 1003), this is the best-supported evolutionary scenario according to our data with *Vainionora* being sister to both of them (Fig. 2A).

**Figure 2.**
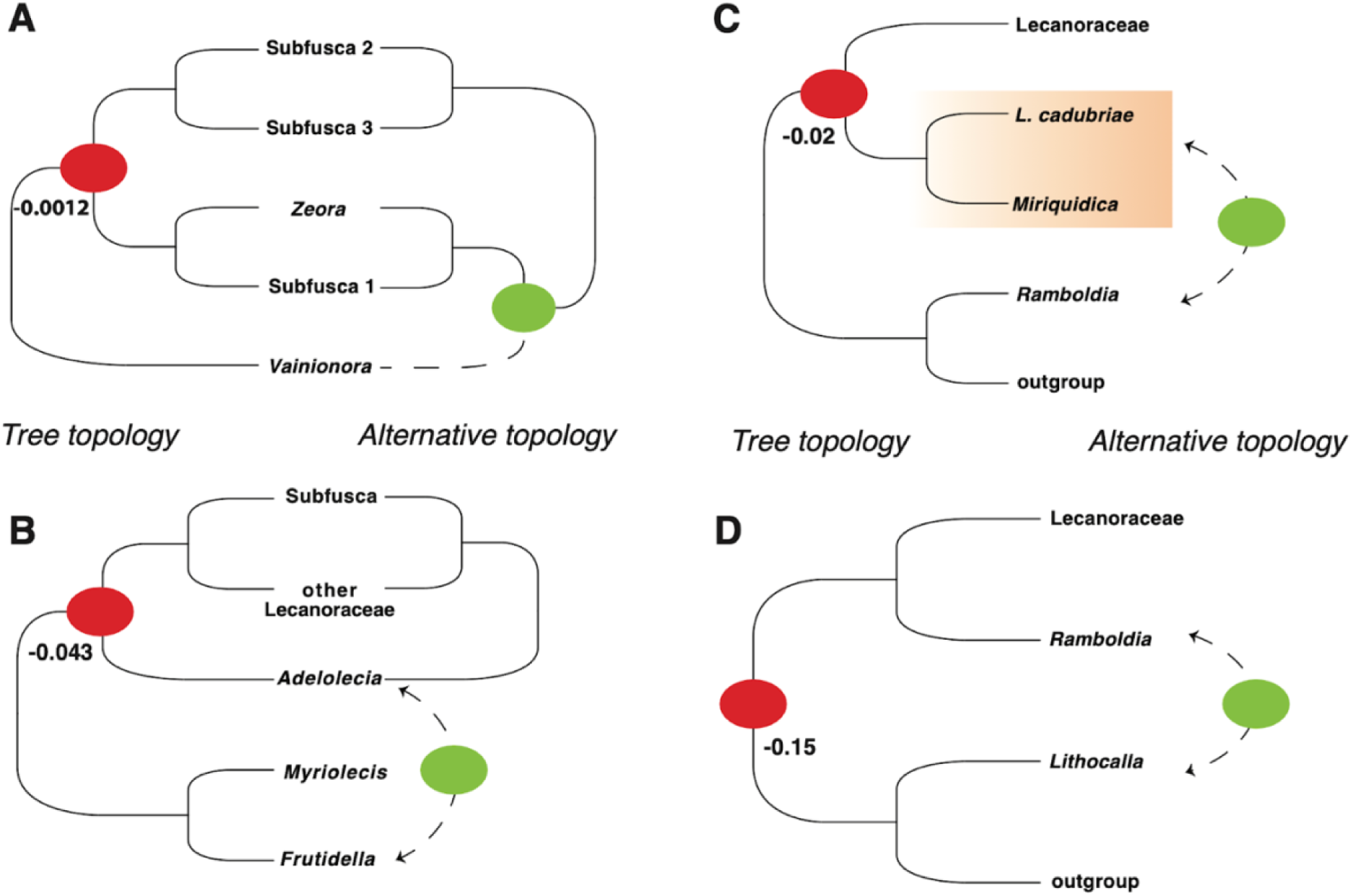
Phylogenetic disagreements with Quartet Concordance (QC) scores for the branches with counter-support. The red circle indicates counter support, with the corresponding QC score displayed next to it. The green circle indicates a supported alternative topology.

Counter-support was also revealed for the node connecting *Adelolecia pilati* to all the species of *Lecanora* s. str., *Glaucomaria* s. lat., and “*Lecidea*” *tuberculifera* (QC/QD/QI: -0.02/0.75/0.48), but with high support (LPP = 0.99). The number of quartet samples supporting this topology was 1740, while the alternative topology suggesting *A. pilati* as sister to a clade containing *Frutidella caesioatra* and *Bryonora* cf. *yeti* was supported by 1904 quartets (Fig. 2B). There were 27 informative concordant genes out of 1003 at this node, which may be explained by the fact that we could extract only 122 orthologous genes for *Adelolecia pilati*.

Two further nodes received counter-support due to uncertainty about the position of *Ramboldia* (QC/QD/QI: -0.15/0.3/0.5), and the *L. fuscescens* group and *Miriquidica* (QC/QD/QI: - 0.02/0.77/0.5). In the ML tree (Fig. 1) these two groups are sisters to *Lecanoraceae* whereas the alternative topologies placed them together with the outgroup, questioning their close relationship with *Lecanoraceae* (Fig. 2C & D). Although we could retrieve only seven orthologous genes for *Lithocalla japonica*, the placement of this species in relation to *Lecanoraceae* remains unaffected in the alternative topologies. All inferred phylogenies placed *Lithocalla* as the sister to *Lecanoraceae* with high support (LPP = 0.99, QC/QD/QI: 0.8/0.22/0.57). The alternative topologies with either *Ramboldia* or the *L. fuscescens* group and *Miriquidica* more closely related to the outgroup, leave *Lithocalla* in this position, based on a high proportion of quartet samples (506). This result aligns with previous studies, which questioned the placement of *Lithocalla* within *Ramalinaceae* (Orange 2020). Thus, it is likely that *Lithocalla* belongs to a different, maybe yet to be described family.

## DISCUSSION

This study presents the first phylogenomic approach for reconstructing the phylogenetic relationships within the family *Lecanoraceae* based on high-throughput data using 1003 orthologous genes. We selected about half of all recognized genera and species groups within this family, sampling key representatives per group and genus. Overall, our results align with previous phylogenies based on Sanger sequences, as well as morphological and chemical data (Zhao *et al*. 2016, Ivanovich *et al*. 2025b), confirming that *Lecanoraceae* is a well-supported monophyletic group and that most genera traditionally classified within this family form well-supported clades. Our results provide improved backbone support for the phylogenetic relationships of these groups. The heterogeneity of *Lecanora* in its broad sense was previously recognized (Choisy 1929, Eigler 1969, Zhao *et al*. 2016, Ivanovich *et al*. 2025b), leading to the segregation of various phenotypically distinct genera. In addition, the inclusion of a growing number of lecideoid genera, previously classified in the artificial genus *Lecidea* s. lat., has contributed to the polyphyletic nature of the remaining species of *Lecanora*. This polyphyly is confirmed in our study, and we discuss the relationships among the different groups below, utilizing phylogenetic, morphological, and chemical evidence.

### *Lecanora* s. str. Fig. 3

Traditionally, species of the *L. subfusca* group (*L. subfusca* 1, 2, and 3), considered to represent the genus in its strict sense, were defined as those related to the type species *L. allophana* (located in *L. subfusca* clade 3) (Brodo & Vitikanen 1984, Vitikanen & Brodo 1985) and characterized by a crustose thallus, lecanorine apothecia, the presence of small or large KOH-insoluble amphithecial crystals, the production of atranorin, and filiform conidia (Brodo 1984; Lumbsch *et al*. 1994). According to our phylogenetic results, the core of *Lecanora* comprises *L. subfusca* clades 1, 2 and 3, *Zeora,* the *Vainionora* group and two more species, *Lecanora masana* and *L. oreinoides*, forming a highly supported clade with strong support (LPP = 100, Phyparts: 884/1003, QC/QD/QI: 0.5/0.82/0.64). In addition, our results reveal that *Zeora* is part of *Lecanora* s. str. being sister to *L. subfusca* clade 1. Similar results were recently presented by Ivanovich *et al*. (2025b), based on a seven-locus phylogeny. Species of *Zeora* are morphologically and chemically distinct from those of the *L. subfusca* group. They are characterized by a yellowish thallus, typically sorediate or farinose in appearance, apothecia with a tendency to become biatorine, the absence of large amphithecial crystals but rather KOH-soluble small crystals, filiform conidia and the production of usnic acid and zeorin instead of atranorin (Eigler 1969, Ivanovich *et al*. 2025b), making *Zeora* a readily recognizable genus. Although the inclusion of *Zeora* in the *L. subfusca* group divides the core of *Lecanora* into three well-supported clades, we were unable to detect any distinguishing characters that would support this separation among the *L. subfusca* group clades. Interestingly, our analyses revealed a relatively high degree of discordance for *Zeora* as a sister to *L. subfusca* clade 1 with only 180 informative concordant genes out of a total of 1003. This indicates a complex evolutionary history of *Lecanora*, which may involve gene flow, rapid radiation, and/or hybridization events. Adding more genes to phylogenetic analyses will not necessarily resolve this issue. Instead, including more taxa from various groups might be more effective to test evidence for any of these specific events, which was outside the scope of this study. In this context, it would also be interesting to date evolutionary events. Because limited sampling is known to strongly affect phylogenetic date estimations (Linder *et al*. 2005), this further underlines the necessity to sample taxa more comprehensively in future studies.

**Figure 3.**
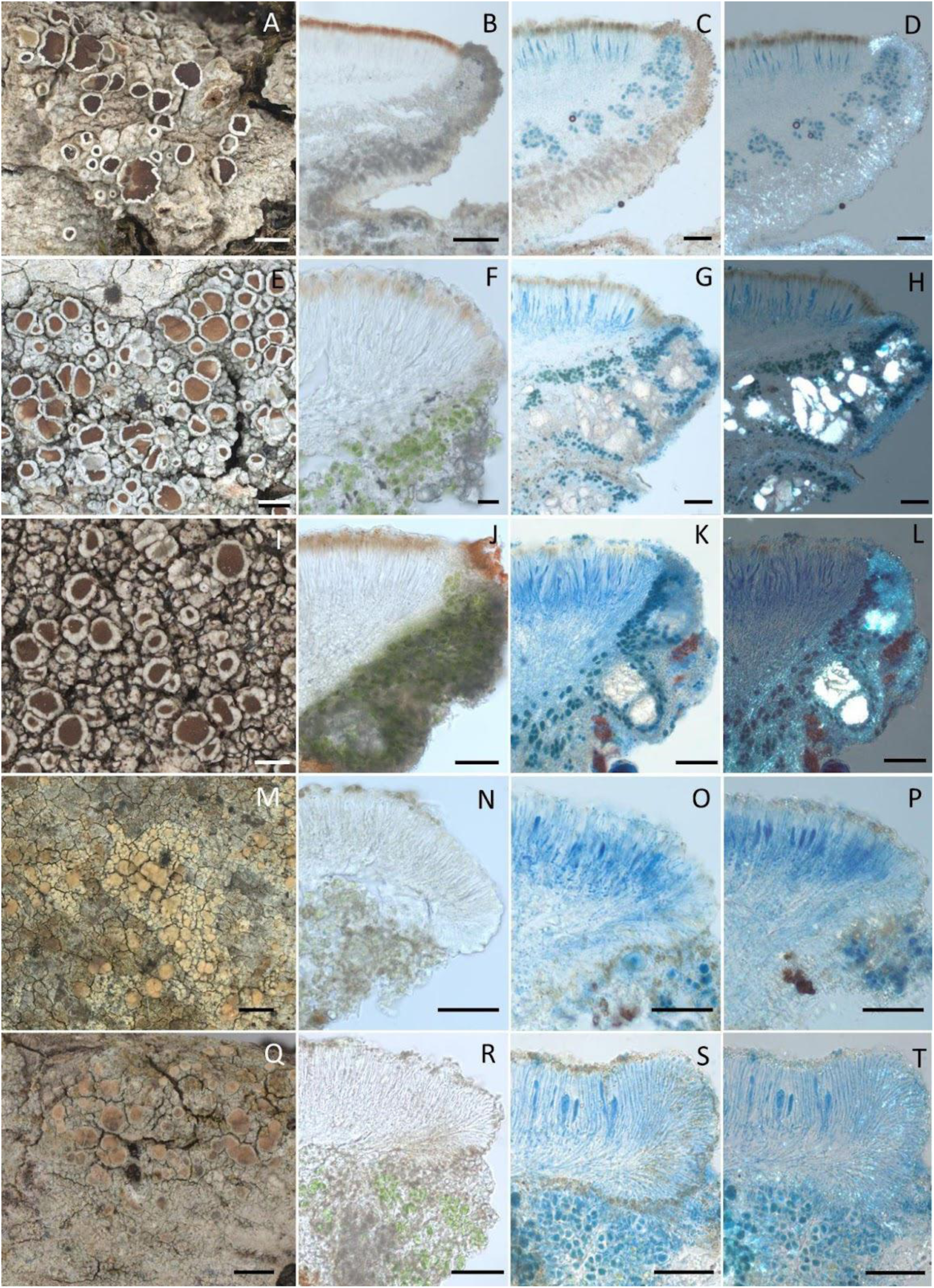
**A-L**, Selected species in the *L. subfusca* group included in our study. **A-D**, *Lecanora allophana* (FR-0264966). A, thallus and apothecia. B, apothecial section in water. C, apothecial section in LCB. D, apothecial section in LCB under polarized light. Scales: A = 1 mm, B, C & D = 100 µm. **E-H**, *Lecanora argentat*a (FR-xxxxxxx). E, thallus and apothecia. F, apothecial section in water. G, apothecial section in LCB. H, apothecial section in LCB under polarized light. Scales: E = 1 mm, F = 20 µm, G & H = 50µm. **I-L**, *Lecanora subimmergens* (NY-3033132). I, thallus and apothecia. J, apothecial section in water. K, apothecial section in LCB. L, apothecial section in LCB under polarized light. Scales: I = 0.5 mm, J, K & L = 50 µm. **M-T**, Selected species in the genus *Zeora* included in our study. **M-P**, *Zeora symmicta* (FR-0265888). M, thallus and apothecia. N, apothecial section in water. O, apothecial section in LCB. P, apothecial section in LCB under polarized light. Scales: M = 1 mm, N, O & P = 50 µm. **Q-T**, *Zeora straminea* (FR-0279047). Q, thallus and apothecia. R, apothecial section in water. S, apothecial section in LCB. T, apothecial section in LCB under polarized light. Scales: Q = 1 mm, R, S & T = 50 µm.

Our results support the idea that the *L. subfusca* group is morphologically coherent but phylogenetically polyphyletic with the inclusion of the recently resurrected genus *Zeora* (Ivanovich *et al*. 2025b). While phylogenomic data improved the support for most internal branches, they also revealed discordance and counter-support at several deeper nodes, keeping the question of the genus boundary open. Given the lack of clear diagnostic traits among the *L. subfusca* group clades and the remaining uncertainty in the backbone topology, we refrain from splitting the group into separate genera. Instead, we propose retaining the concept of the *L. subfusca* group for the time being to reflect its phenotypic unity, while acknowledging its polyphyly.

### The “*Vainionora* group” Fig. 4

The genus *Vainionora* is represented in our phylogeny by a single species, *Lecanora markjohnstonii*. This species was originally described as a sterile *Lecanora* from North America (Anderson Stewart *et al*. 2018). However, Ivanovich *et al*. (2025b) recently discovered that the Sanger sequences used in their study were identical to those from fertile material collected in Rwanda. This finding highlighted characteristic traits of *Vainionora*, such as a pigmented hypothecium and bacilliform conidia (Kalb 1991, Bungartz *et al*. 2020, Weber *et al*. in rev.), showing a close relationship with *Lecanora alboflavida* and *Lecanora orientoafricana*. These latter taxa also show characters typical for *Vainionora* (Ivanovich *et al*. 2025b). In our phylogeny, it formed a sister clade to *L. masana* and *L. oreinoides* with strong support (LPP = 100, Phyparts: 712/1003, QC/QD/QI: 0.4/0.8/0.57). As outlined above, our phylogenetic results revealed the “*Vainionora* group”, which includes *L. markjohnstonii*, *L. masana*, and *L. oreinoides*, as a sister clade to the three *L. subfusca* groups and *Zeora* (Fig. 1). In the alternative supported topology, *Vainionora* is sister to *L. subfusca* clade 1 and *Zeora* (Fig. 2A). Morphologically, *L. masana* fits within the *L. subfusca* group due to its hyaline hypothecium and filiform conidia (Lendemer *et al*. 2013). Contrarily, *Lecanora oreinoides* is morphologically highly distinctive, characterised by immersed apothecia, the absence of amphithecial crystals, and bacilliform conidia. The well-supported close relationship among this species, *Vainionora* and *Zeora* with typical *Lecanora* provides further evidence that phenotypical and molecular evolution are not aligned in *Lecanora* s. str.

**Figure 4.**
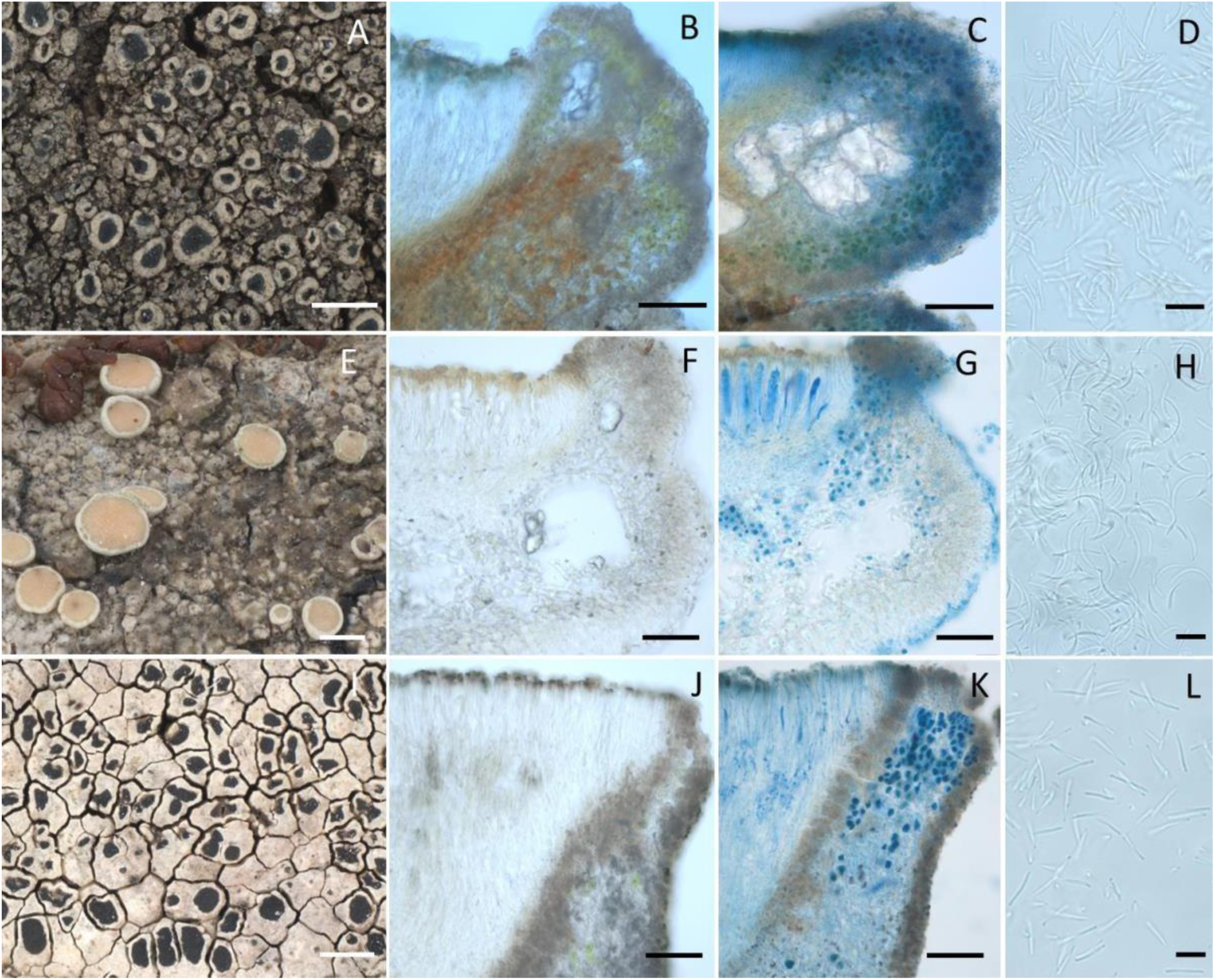
**A-L**, Selected species in the “*Vainionora* group” included in this study. **A-D**, “*Vainionora” markjohnstonii* (FR-0175442). A, thallus and apothecia, scale = 1 mm. B, apothecial section in water. C, apothecial in LCB. D, bacilliform conidia. Scales: A = 1 mm, B & C = 50 µm, D = 10 µm. **E-H**, *Lecanora masana* (NY-2606763). E, thallus and apothecia. F, apothecial section in water. G, apothecial section in LCB. H, filiform conidia. Scales: E = 0.5 mm, F & G = 50 µm, H = 10µm. **I-L**, *Lecanora subimmergens* (NY-3033132). I, thallus and apothecia. J, apothecial section in water. K, apothecial section in LCB. L, bacilliform conidia. Scales: I = 1 mm, J & K = 50 µm, L = 10 µm.

### “*Glaucomaria* s. lat.” Fig. 5

Our phylogenomic results clarified the positions of four lineages that were traditionally classified within *Lecanora*: *Glaucomaria*, *Verseghya*, and the *L. intumescens* and *L. albella* groups (Imshaug & Brodo 1966, Leuckert & Poelt 1989, Lumbsch *et al*. 1997, Grube 2004, Kondratyuk *et al*. 2016, Brodo *et al*. 2019). These groups formed a highly supported clade (LPP = 100, Phyparts: 885/1003, QC/QD/QI: 0.27/0.38/0.5). Although these lineages share certain phenotypic traits with *Lecanora* s. str., such as lecanorine apothecia with small amphithecial crystals, the presence of atranorin, and filiform conidia, they also exhibit differences in the development of the thalline margin, frequent presence of apothecial pruina, and secondary metabolites (Imshaug & Brodo 1966, Poelt 1989, Lumbsch *et al*. 1997, Brodo *et al*. 2019, Arup *et al*. 2023).

**Figure 5.**
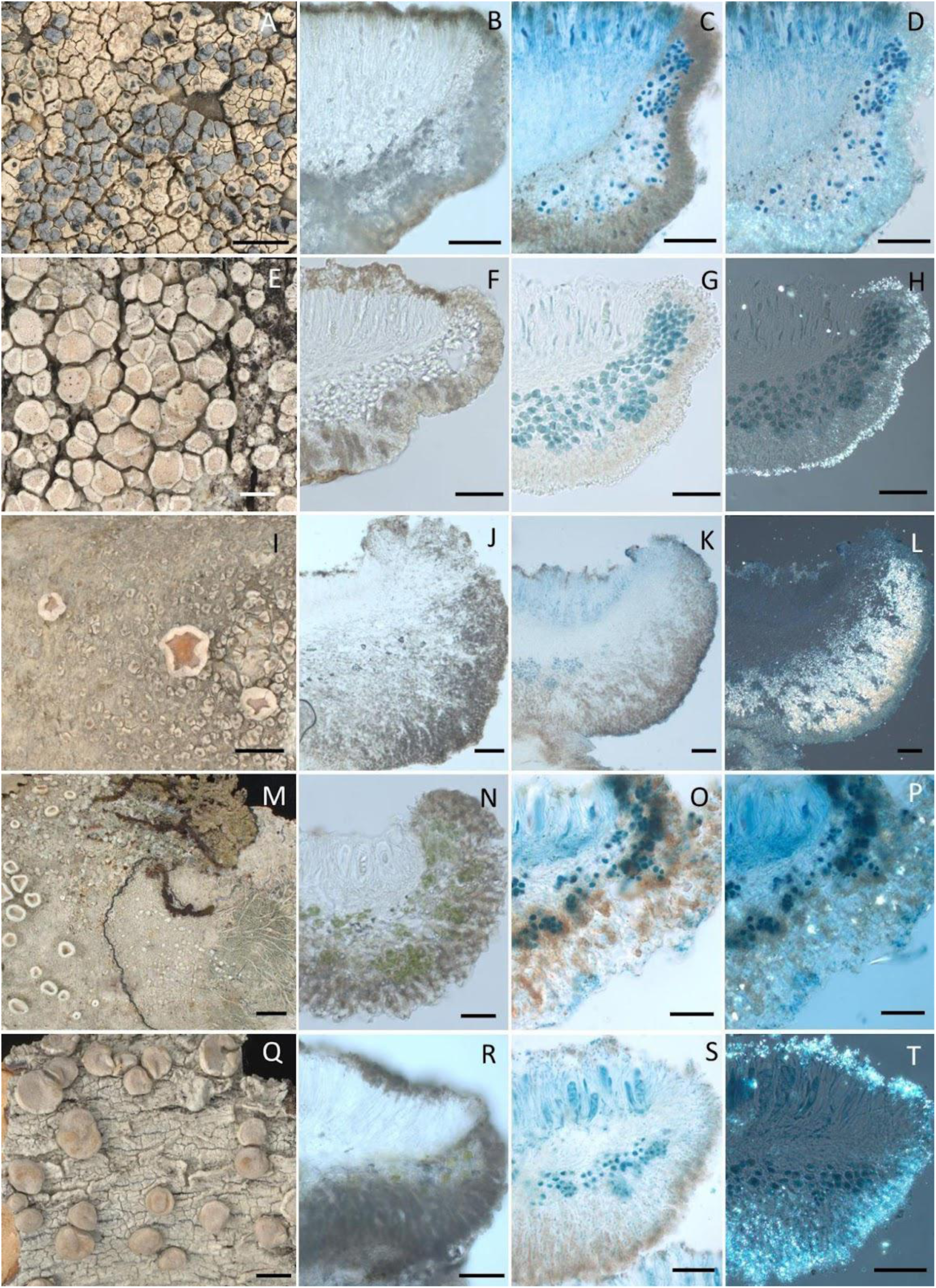
**A-T**, Selected species from “*Glaucomaria* s.lat.” included in this study. **A-D**, *Glaucomaria bicincta* (FR-0175453). A, thallus and apothecia, scale = 1 mm. B, apothecial section in water. C, apothecial in LCB. D, apothecial section in LCB under polarized light. Scales: A = 2 mm, B, C & D = 50 µm. **E-H**, *Glaucomaria carpinea* (FR-0264946). E, thallus and apothecia. F, apothecial section in water. G, apothecial section in LCB. H, apothecial section in LCB under polarized light. Scales: E = 0.5 mm, F, G & H = 50 µm. **I-L**, *Lecanora intumescens* (FR-0279044). I, thallus and apothecia. J, apothecial section in water. K, apothecial section in LCB. L, apothecial section in LCB under polarized light. Scales: I = 2 mm, J, K & L = 100 µm. **M-P**, *Verseghya thysanophora* (FR-0265868). M, thallus and apothecia. N, apothecial section in water. O, apothecial section in LCB. P, apothecial section in LCB under polarized light. Scales: M = 1 mm, N, O & P = 50 µm. **Q-T**, *Lecanora albella* (FR-xxxxxxx). Q, thallus and apothecia. R, apothecial section in water. S, apothecial section in LCB. T, apothecial section in LCB under polarized light. Scales: Q = 1 mm, R, S & T = 50 µm.

According to our phylogeny, the clade including these groups is closely related to *Lecanora* s. str. *Glaucomaria* (Choisy 1929, Ivanovich *et al*. 2025b) is shown to be sister to clades composed of *L. intumescens*, the recently described genus *Verseghya,* and *L. albella*. Each of these four clades is morphologically distinct and easily distinguishable from one another. *Glaucomaria* is characterized by a well-developed amphithecial cortex, heavily pruinose apothecia, and the production of sordidone. The species in the *L. intumescens* group are defined by a thick, prominent, typically flexuose, whitish apothecial margin, an amphithecial pseudocortex, commonly epruinose apothecia, and the production of zeorin. *Verseghya* includes sterile or sorediate species with a distinctive fibrous prothallus, producing both zeorin and usnic acid. Finally, the *L. albella* group comprises species with heavily pruinose apothecia, an amphithecial pseudocortex, and β-orcinol depsidones. The tendency to form pruinose apothecia, the absence of large amphithecial crystals, atranorin as the primary secondary compound, and the presence of filiform conidia all support the inferred shared ancestry of these groups.

The systematic placement of *Verseghya* has been uncertain, as its position within the ’*Verseghya*–*Lecidella*–*Pyrrhospora*’ clade received poor support (Kondratyuk *et al*. 2019). Our phylogenomic analysis provides strong evidence for a close relationship between *Verseghya* and morphologically similar lecanorine genera. Previous phylogenetic studies (e.g., Zhao *et al*. 2016, Brodo *et al*. 2019, Ivanovich *et al*. 2025b) recognized the *L. intumescens* and *L. albella* groups as independent lineages, but neither group has been formally classified as separate genera. A more comprehensive taxon sampling is necessary to determine whether these groups should merit generic rank or if they should be included into *Glaucomaria*.

### “MPRPS clade” Fig. 6

The so-called “MPRPS clade” was first introduced by Medeiros *et al*. (2021) to describe a well-supported lineage that includes the genera *Myriolecis* (*L. dispersa* group), *Protoparmeliopsis* (*L. muralis* group), *Rhizoplaca*, *Carbonea*, *Straminella* (*L. varia* group), *Lecanoropsis* (*L. saligna* group), and *Bryonora* (Zhao *et al*. 2016, Medeiros *et al*. 2021, Svensson *et al*. 2021, Zhang *et al*. 2022, Ivanovich *et al*. 2025a).

**Figure 6.**
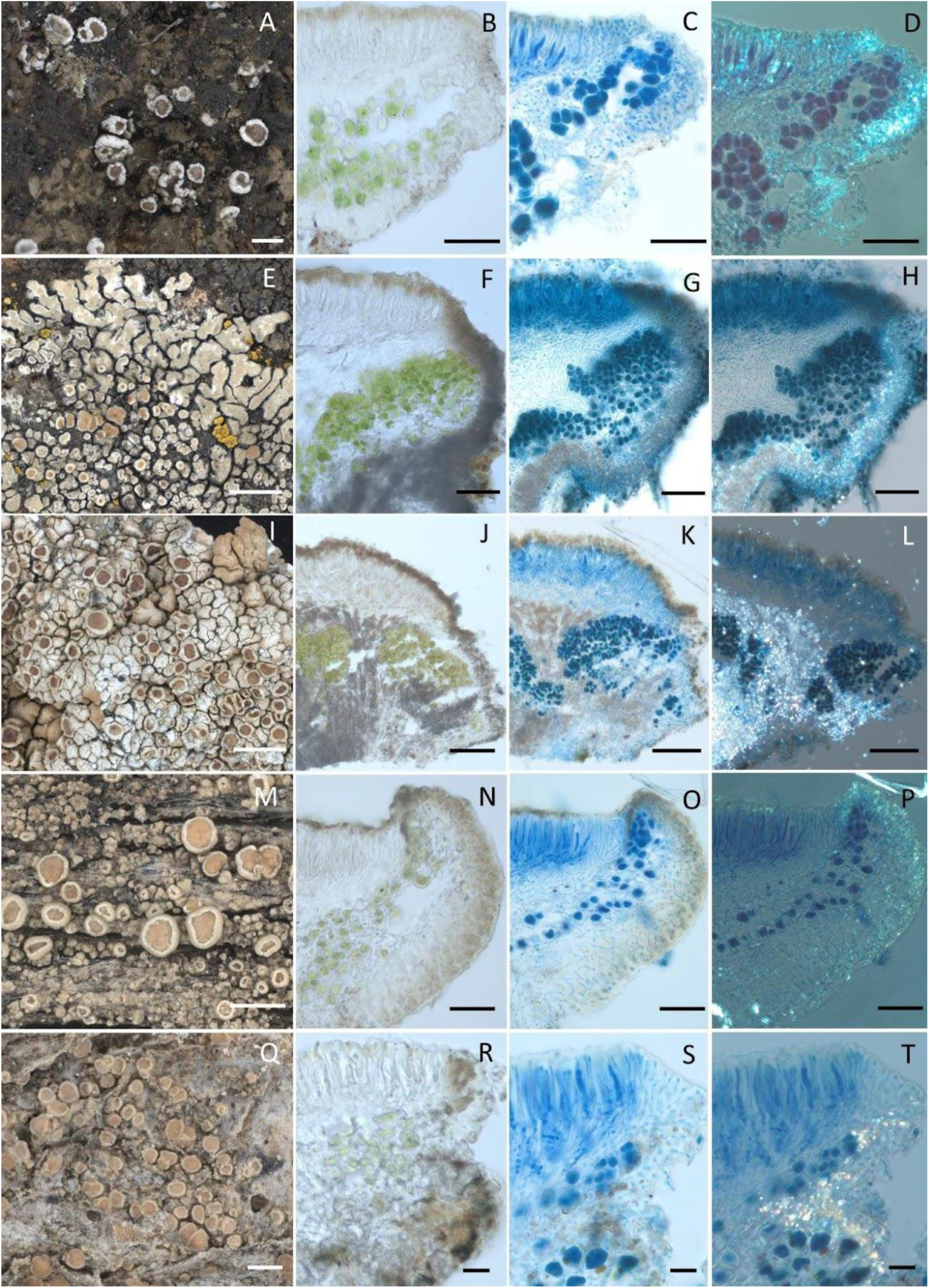
**A-T**, Selected species in the “MPRPS” clade included in this study. **A-D**, *Myriolecis dispersa* (FR-0264963). A, thallus and apothecia. B, apothecial section in water. C, apothecial in LCB. D, apothecial section in LCB under polarized light. Scales: A = 1 mm, B, C & D = 20 µm. **E-H**, *Protoparmeliopsis muralis* (FR-0264962). E, thallus and apothecia. F, apothecial section in water. G, apothecial section in LCB. H, apothecial section in LCB under polarized light. Scales: E = 2 mm, F, G & H = 50 µm. **I-L**, *Rhizoplaca pachyphylla* (FR-xxxxxxx). I, thallus and apothecia. J, apothecial section in water. K, apothecial section in LCB. L, apothecial section in LCB under polarized light. Scales: I = 2 mm, J, K & L = 100 µm. **M-P**, *Straminella varia* (FR-0261120). M, thallus and apothecia. N, apothecial section in water. O, apothecial section in LCB. P, apothecial section in LCB under polarized light. Scales: M = 1 mm, N, O & P = 50 µm. **Q-T**, *Lecanoropsis subsaligna* (FR-0279046). Q, thallus and apothecia. R, apothecial section in water. S, apothecial section in LCB. T, apothecial section in LCB under polarized light. Scales: Q = 0.5 mm, R, S & T = 20 µm.

Subsequent phylogenetic studies confirmed the close relationship within this clade. With the exception of *Carbonea*, these groups were previously classified under *Lecanora* s. lat. Although there is consistent support for the clade, the phylogenetic relationships among its members, and the placement of *Bryonora* and *Carbonea*, have varied in different studies.

Our data further clarifies the relationships within the MPRPS clade. Consistent with previous findings, our phylogenetic results show that the MPRPS group forms a clade with maximum support (LPP = 100, Phyparts: 997/1003, QC/QD/QI: 0.9/1/0.9). Notably, while the lecidoid genus *Carbonea* is included within this clade, *Bryonora* is excluded, showing instead a close relationship to *Frutidella*. Our results provide strong evidence for a well-supported sister relationship between *Myriolecis* and *Protoparmeliopsis* (LPP = 100, Phyparts: 681/1003, QC/QD/QI: 0.22/0.33/0.54), which aligns with several previous classifications (Zhao *et al*. 2016; Zhang *et al*. 2022). This group is sister to a clade comprising *Rhizoplaca* and *Carbonea* (LPP = 98). However, the high discordance of tree topologies indicates considerable uncertainty at this node with counter-support (Phyparts: 256/1003, QC/QD/QI: -0.043/0.63/0.46). An alternative supported topology suggests that *Carbonea* might be sister to *Straminella*. Previous phylogenetic studies showed *Carbonea* being a part of either the *L. polytropa* group (Zhang *et al*. 2022) or *Rhizoplaca* (Svensson & Westberg 2021), suggesting a potential close relationship between the respective groups. Zhao *et al*. (2016) included all three groups in their two-locus phylogenetic analysis, but the results did not support a close relationship among these lineages. Since a representative of the *L. polytropa* group could not be included in our dataset, the position of *Carbonea* within the MPRPS clade remains unclear.

Additionally, our results strongly support the recognition of *Straminella* as a distinct genus, separate from *Zeora* or *Lecanoropsis*. Ivanovich *et al*. (2025a, b) recently resurrected these three genera based on morphological and chemical traits. Although the backbone of the seven-locus phylogeny provided by these authors was relatively poorly supported, our phylogenomic data strongly supports a sister relationship between *Straminella* and the clade formed by *Myriolecis*, *Protoparmeliopsis*, *Rhizoplaca*, and *Carbonea* (LPP = 100, Phyparts: 679/1003, QC/QD/QI: 0.2/0.99/0.52). Meanwhile, *Lecanoropsis* is recovered as the earliest-diverging lineage within the MPRPS clade (LPP = 100, Phyparts: 997/1003, QC/QD/QI: 0.9/1/0.9).

The genus *Bryonora* was segregated from *Lecanora* by Poelt (1983), but its phylogenetic position remained uncertain, with some studies suggesting a close relationship to *Protoparmelia* or other genera that are no longer included in *Lecanoraceae* (Poelt & Obermayer 1991, Grube *et al*. 2004). However, subsequent multigene phylogenetic analyses based on Sanger sequences confirmed its inclusion in *Lecanoraceae* (Zhao *et al*. 2016). Medeiros *et al*. (2021) considered *Bryonora* to be a part of the MPRPS clade, but this affiliation has later been challenged by several studies (Svensson *et al*. 2022, Ivanovich *et al*. 2025b). Our phylogenetic analyses instead recovered *Bryonora* as the sister clade to *Frutidella* (LPP = 100, Phyparts: 11/1003, QC/QD/QI: 0.65/0.43/0.66), one of the lecideoid genera within *Lecanoraceae*, positioning it at the deep node as the sister to all other clades of the family *Lecanoraceae*. Even though we could recover only 11 orthologous genes for *Bryonora* due to insufficient sequencing depth, all gene topologies supported this relationship.

### Lecideoid genera within *Lecanoraceae* Fig. 7 & Fig. 8

In Zahlbruckner’s (1926) classification, genera assigned to *Lecanoraceae* were characterized by lecanorine apothecia, while similar taxa with a proper apothecial margin were typically placed in *Lecideaceae*. Hafellner (1984) modified our understanding of the family *Lecanoraceae* by moving the lecideoid genus *Lecidella* to it based on ascus apical structures. *Miriquidica,* which comprises species with lecanorine and lecideine apothecia, was originally described within *Lecanoraceae* as closely related to *Bryonora* and *Protoparmelia* (Hertel & Rambold 1987).

**Figure 7.**
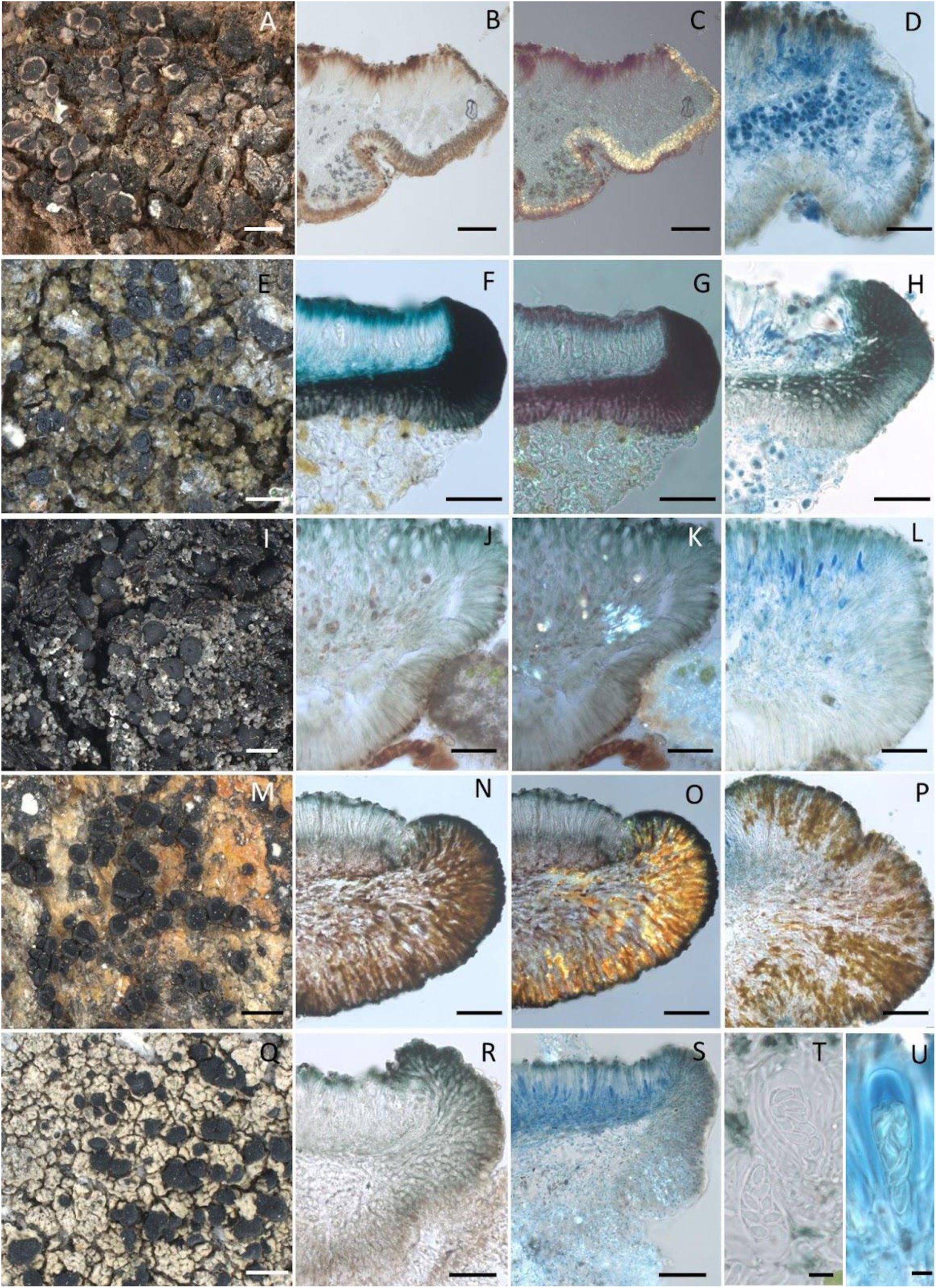
**A-D**, *Bryonora* cf. *yeti* (FR-xxxxxxx). A, thallus and apothecia. B, apothecial section in water. C, apothecial section under polarized light. D, apothecial section in LCB. Scales: E = 1 mm, F & G = 100 µm, H = 50 µm. **E-U**, selected lecideoid species in *Lecanoraceae*. **E-H**, *Carbonea vorticosa* (FR-0279047). E, thallus and apothecia. F, apothecial section in water. G, apothecial section under polarized light. H, apothecial section in LCB. Scales: A = 0.5 mm, B, C & D = 50 µm. **I-L**, *Frutidella caesioatra* (FR-xxxxxxx). I, thallus and apothecia. J, apothecial section in water. K, apothecial section under polarized light. L, apothecial section in LCB. Scales: I = 1 mm, J, K & L = 50 µm. **M-P**, *Adelolecia pilati* (O-L-235742). M, thallus and apothecia. N, apothecial section in water. O, apothecial section under polarized light. P, apothecial section. Scales: M = 1 mm, N, O & P = 50µm. **Q-U**, “*Lecidea*” *tuberculifera* (O-L-225040). Q, thallus and apothecia. R, apothecial section in water. S, apothecial section in LCB under polarized light. T, 8-spored asci. U, *Lecanora*-type asci in Lugol’s iodine. Scales: Q = 2 mm, R & S= 50 µm, T & U = 10 µm.

**Figure 8.**
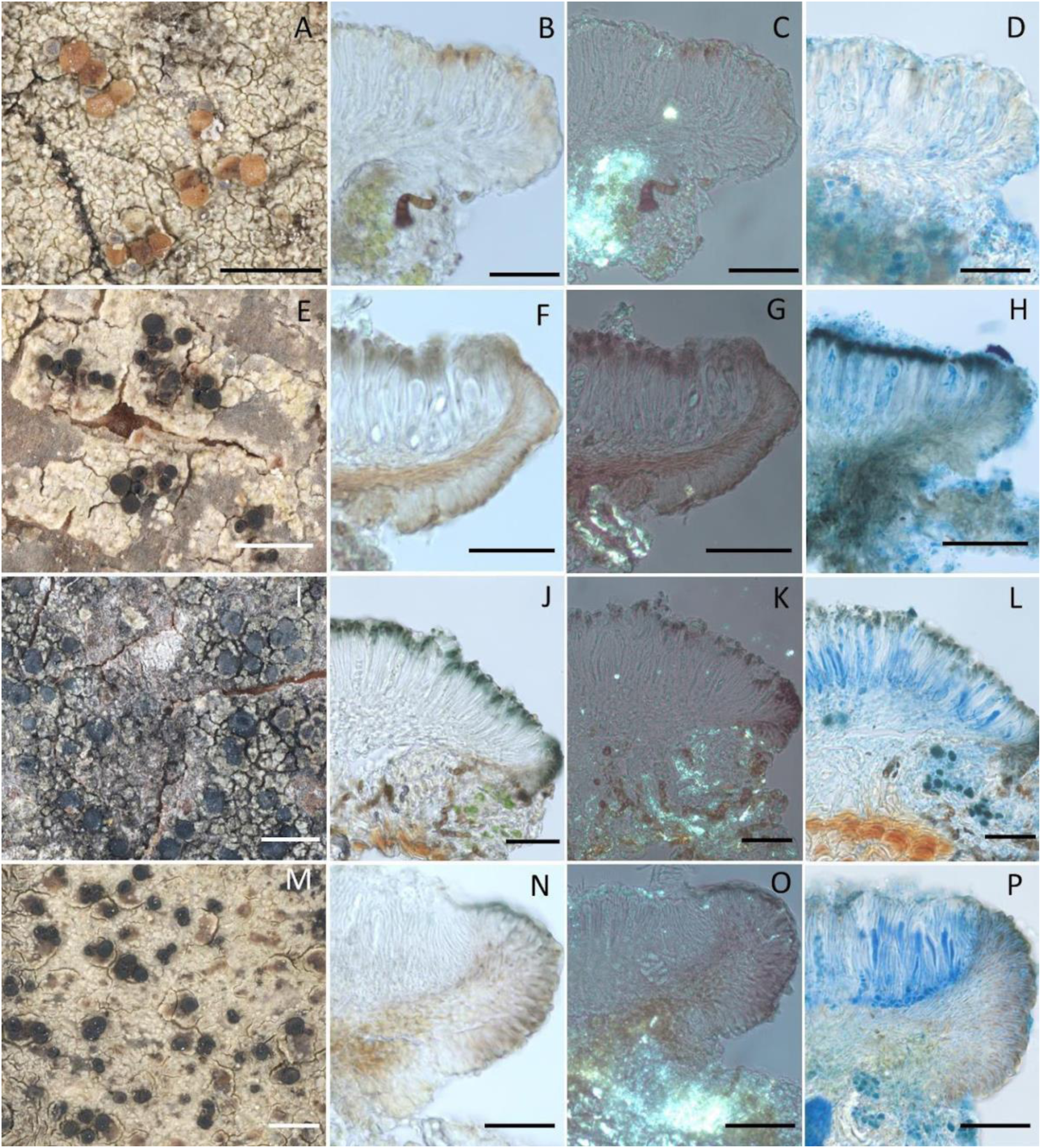
**A-P**, species of the ‘*Lecidella* s. l.’ clade included in this study. **A-D**, *Xanthosyne varians* (NY-3033725). A, thallus and apothecia. B, apothecial section in water. C, apothecial section under polarized light. D, apothecial section in LCB. Scales: A = 1 mm, B, C & D = 50 µm. **E-H**, *Lecidella* sp. (NY-2606722). E, thallus and apothecia. F, apothecial section in water. G, apothecial section under polarized light. H, apothecial section in LCB. Scales: E = 1 mm, F, G & H = 50 µm. **I-L**, *Nimisora iberica* (Pérez-Ortega 12120). I, thallus and apothecia. J, apothecial section in water. K, apothecial section under polarized light. L, apothecial section in LCB. Scales: I = 1 mm, J, K & L = 50 µm. **M-P**, *Lecidella elaeochroma* (FR-0261123). M, thallus and apothecia. N, apothecial section in water. O, apothecial under polarized light. P, apothecial section in LCB. Scales: M = 1 mm, N, O & P = 20 µm.

Recent phylogenetic studies supported the inclusion of further lecideoid genera, such as *Adelolecia*, *Frutidella*, *Japewia*, *Nimisora*, *Palicella*, *Pyrrhospora*, and *Xanthosyne*, into *Lecanoraceae*. These findings underlined that the lecanorine margin is not a defining characteristic of *Lecanoraceae* (Schmull *et al*. 2011, Miadlikowska *et al*. 2014, Rodriguez-Flakus & Printzen 2014, Zhao *et al*. 2016, Kistenich *et al*. 2018). Moreover, taxa with a proper margin seem to be concentrated around the deeper nodes of the *Lecanoraceae*, indicating that the lecanorine margin might be a derived character. *Adelolecia* was historically classified under *Bacidiaceae* (Hafellner 1984) and was later moved to *Ramalinaceae* (Hertel & Rambold 1995). However, subsequent molecular studies supported its position within *Lecanoraceae* (Ekman *et al*. 2008, Zhao *et al*. 2016, Kistenich *et al*. 2018, Kondratyuk *et al*. 2019), although its relation to other genera remained unclear. Our phylogeny shows its sister position to most of *Lecanora* s. lat. and “*Lecidea*” *tuberculifera* (Fig. 1). However, only 27 concordant gene trees supported this placement, while an alternative topology suggests its sister relationship to *Frutidella* and *Bryonora* (Fig. 2B). Consequently, the phylogenetic position of the genus remains uncertain and requires further study.

The placement of *Frutidella* within *Lecanoraceae*, rather than *Ramalinaceae*, has already been confirmed by Schmull *et al*. (2011) and Miadlikowska *et al*. (2014). However, its specific phylogenetic position within the family has remained uncertain. Our results demonstrate that *Frutidella* and *Bryonora* are sister lineages, closely related to all other clades of the *Lecanoraceae*. Furthermore, “*Lecidea*” *tuberculifera* seems to be closely related to “*Glaucomaria* s. lat.”. Its distinctiveness from this genus is supported not only by its lecideine apothecia but also by its *Lecidella*-type asci. Given these differences and its phylogenetic position it likely deserves generic rank.

*Lecidella*, the recently described *Nimisora* (Pérez-Ortega *et al*. 2023), and *Xanthosyne* (Brodo *et al*. 2024) form a strongly supported clade (LPP = 100, Phyparts: 989/1003, QC/QD/QI: 0.96/0.8/0.96). These three genera share several morphological characteristics, such as lecideine apothecia, *Lecidella*-type asci, and only weakly conglutinate paraphyses (Zhao *et al*. 2015, Pérez-Ortega *et al*. 2023, Brodo *et al*. 2024). Several previous studies showed *Lecidella* to be sister to *Lecanora* s. lat. and the MPRPS-clade (Zhao *et al*. 2016, Medeiros *et al*. 2021, Brodo *et al*. 2024, Ivanovich *et al*. 2025a). However, our results show that *Lecidella* is instead sister to *Lecanora*, *Glaucomaria* s. lat., and the MPRPS clade (LPP = 100, Phyparts: 410/1003, QC/QD/QI: 0.1/0.67/0.53).

In spite of its similarity with *Lecidella*, *Nimisora* seems to be more closely related to *Ramboldia*, forming a sister clade in the three-gene phylogeny by Pérez-Ortega *et al*. (2023). *Xanthosyne* also shares morphological and chemical characters with *Lecidella,* such as weakly conglutinated paraphyses, *Lecidella*-type asci, and the production of xanthones and atranorin (Brodo *et al*. 2024). In our analysis *Lecidella elaeochroma* formed a separate clade sister to *Nimisora*, *Xanthosyne*, and *Lecidella* sp. from GenBank. Already Zhao *et al*. (2016) demonstrated that *Lecidella* is subdivided into three distinct lineages. However, our taxon sampling representing this clade is still too sparse and the identification of the *Lecidella* sp. specimen has to be confirmed upon final taxonomic decision. Furthermore, additional lecideoid genera, such as *Traponora*, *Palicella*, and *Japewiella*, are morphologically similar to these lineages and should be included in a phylogenomic analysis before any taxonomic conclusions are drawn. For this reason, we leave it open to discussion whether these three genera should be combined or if *Lecidella* should be split into several genera.

### The clades sister to Lecanoraceae: L. fuscescens group, Ramboldia, and Miriquidica Fig. 9

*Miriquidica* was initially classified in *Lecanoraceae*, but recent studies showed its placement outside of this family (Miadlikowska *et al*. 2014, Svensson *et al*. 2022). When *Miriquidica* was again included within *Lecanoraceae*, its phylogenetic relationship with other members of the family remained unclear due to poor phylogenetic support (e.g., Zhao *et al*. 2016, Ivanovich *et al*. 2025b). Our results show that *Miriquidica* forms a distinct clade sister to *Lecanoraceae* and closely related to the *L. fuscescens* group. Indirectly, this seems to corroborate the well-supported relationship between *L. physciella* and *Miriquidica* demonstrated by Zhao *et al*. (2016). The seven-locus phylogeny by Ivanovich *et al*. (2025b) showed *L. physciella* nested with *L. cadubriae*, although this finding lacks strong support. However, since *L. physciella* is a saxicolous, Antarctic endemic that produces usnic acid (Sliwa & Olech 2002), it differs significantly from *L. cadubriae*, which occur on wood and bark in the Northern Hemisphere and produce norstictic acid and/or connorstictic acid and salazinic acid. For this reason, we hypothesise that this relationship may be an artefact resulting from limited taxon sampling in Ivanovich *et al*. (2025b).

**Figure 9.**
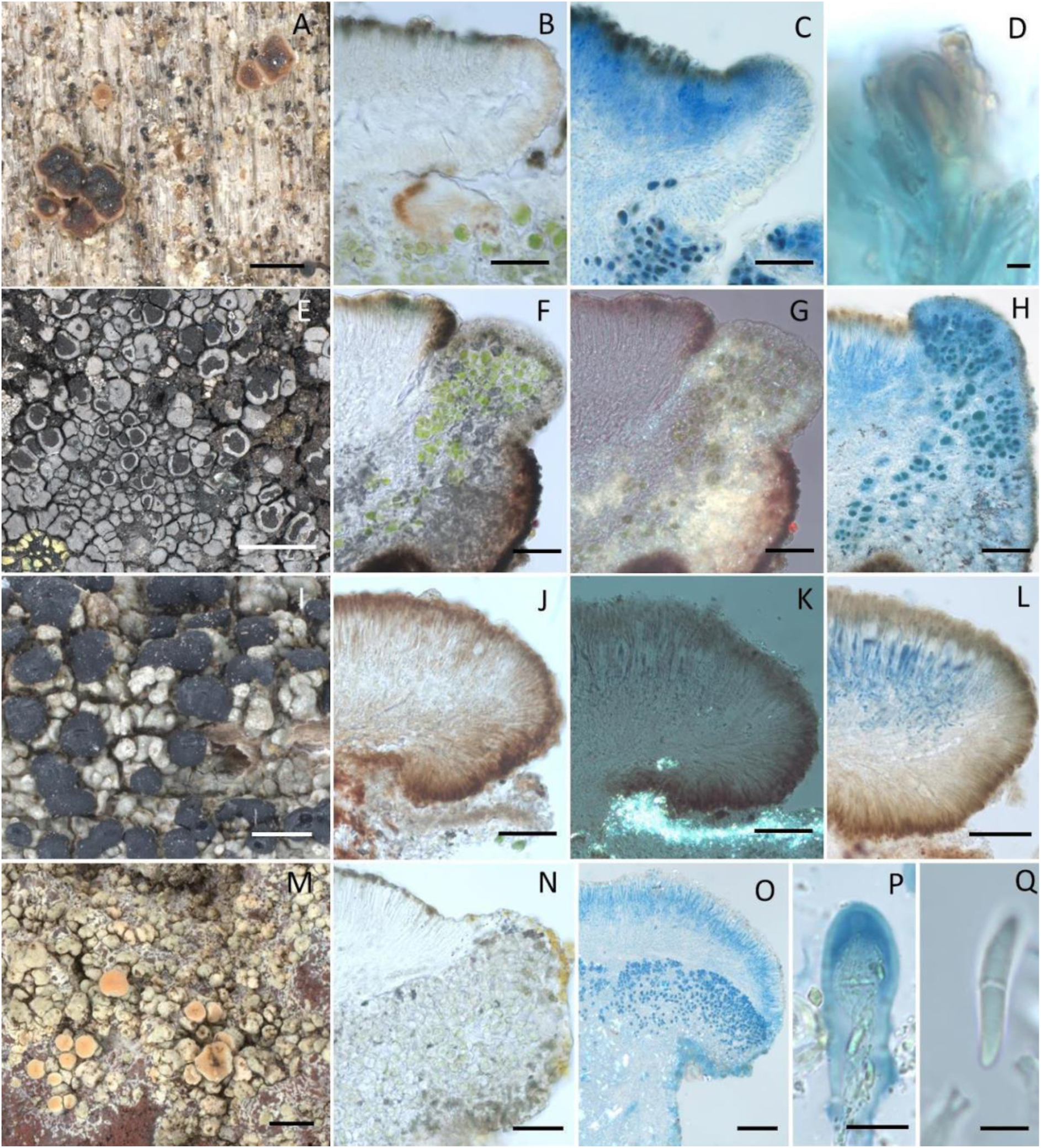
**A-D**, *Lecanora cadubriae* (FR-0279001). A, thallus and apothecia. B, apothecial section in water. C, apothecial in LCB. D, ascus in Lugol’s iodine. Scales: A = 0.5 mm, B & C = 50 µm, D = 10 µm. **E-H**, *Miriquidica complanata* (O-L-227468). E, thallus and apothecia. F, apothecial section in water. G, apothecial section in water under polarized light. H, apothecial section in LCB. Scales: E = 2 mm, F, G & H = 50 µm. **I-L**, *Ramboldia elabens* (FR-0264965). I, thallus and apothecia. J, apothecial section in water. K, apothecial section in water under polarized light. L, apothecial section in LCB. Scales: I = 1 mm, J, K & L = 20 µm. **M-P**, *Lithocalla japonica* (FR-0265882). M, thallus and apothecia. N, apothecial section in water. O, apothecial section in LCB under polarized light. P, ascus in Lugol’s iodine. Q, ascospore. Scales: M = 1 mm, N = 50 µm, O = 100 µm, P = 10 µm, Q = 5 µm.

Previous phylogenetic studies showed *Ramboldia* as a well-supported clade sister to *Lecanoraceae* (Miadlikowska *et al*., 2014, Zhao *et al*., 2016), leading to its inclusion in a separate family, *Ramboldiaceae*. However, this classification has not been unanimously accepted, as *Ramboldia* has also been treated as part of *Lecanoraceae* in both morphological and phylogenetic studies (Kantvilas 2016, Zhao *et al*. 2016, Elix & McCarthy 2017, Medeiros *et al*. 2021, Ivanovich *et al*. 2025b). Our phylogenetic results confirm that *Ramboldia* forms a clade sister to the remaining *Lecanoraceae*. However, the alternative topologies inferred by quartet sampling placed *Miriquidica* and *Ramboldia* as sisters to the outgroup, questioning their close relationship with *Lecanoraceae* (Fig. 2C & D).

### Lithocalla Fig. 9

The genus *Lithocalla* was recently described within the family *Ramalinaceae*, accommodating two sterile species. Its phylogenetic relationship to the family *Ramalinaceae* remained somewhat uncertain due to poor support (Orange 2020). The genus was only included in our study because the sample, an undescribed fertile species of the genus collected in Japan, was misidentified as a member of the *L. polytropa* group due to its morphological similarity with members of this group, and was subsequently found through sequence comparison to be more closely allied with *Lithocalla*. To clarify its systematic placement, we included this species in our studies. Our results suggest that *Lithocalla* may represent an independent lineage, distinct from both *Ramalinaceae* and *Lecanoraceae*. However, this conclusion should not be over-interpreted as only seven loci were available for phylogenomic reconstruction.

## CONCLUSIONS

The phylogenomic analysis of the *Lecanoraceae* family reveals complex evolutionary relationships among its genera, illustrating that simply adding more genetic loci does not always clarify uncertainties in deeper phylogenetic nodes. Various processes, such as incomplete lineage sorting, gene duplication, gene loss, introgressive hybridization, and horizontal gene transfer, contribute to the mosaic nature of genomes and can lead to discordance in gene trees (Maddison 1997, Mallet *et al*. 2016, Arnold & Kunte 2017, Walker *et al*. 2024). Furthermore, limited sampling has been shown to significantly impact phylogenetic dating (Linder *et al*. 2005).

To effectively address these complexities, future studies should focus on including a broader range of taxa with the inclusion of a higher number of genes. For example, the *Lecanora subfusca* group has shown inconsistencies across phylogenetic analyses, raising questions about its monophyly and the delimitation of the genus in a strict sense. The core group of *Lecanoraceae* contains all taxa with the distinctive “*Lecanora* phenotype,” defined by the presence of apothecia with a thalline margin, containing crystalline deposits, and the production of atranorin. Within this core group, we recognize the paraphyletic *L. subfusca* group (*Lecanora* s. str.), *Vainionora*, and *Glaucomaria* s. lat. Notably, the genus *Zeora*, despite its morphological and chemical divergence from typical *Lecanora* species, nested within *Lecanora* s. str. Species of *Zeora* display distinct characteristics in having either lecanorine apothecia with margins lacking a clearly defined cortex and oxalate crystals, or in having biatorine apothecia, producing usnic acid and zeorin instead of atranorin as a major compound.

Additionally, the MPRPS clade and the genus *Lecidella* formed a separate clade outside the core *Lecanora* group. The accurate delimitation of the *Lecanoraceae* and the phylogenetic relationship among *Bryonora*, *Frutidella*, *Miriquidica*, *Ramboldia*, and *Lithocalla* remains under question. This uncertainty derives partly from the limited sampling of taxa representing these clades, but perhaps also from the complex evolutionary history, which can be addressed by comparative genome studies of the whole genome architecture.

This underlines the central theme of our inquiry: increasing genetic loci alone does not automatically lead to stronger phylogenetic support at deeper nodes. It is crucial to integrate comprehensive taxon sampling with careful locus selection to clarify the evolutionary history of the *Lecanoraceae* family and enhance our overall understanding of the evolution of lichenized fungi. Despite the complex relationships within the family, our study contributes valuable insights into the phylogenetic relationship among the core taxa within the family *Lecanoraceae* and represents a pioneering effort towards improving our understanding of these relationships based on a large number of orthologous genes.

## ACKNOWLEDGEMENTS

Phylogenetic analyses were performed on the TBG cluster (Senckenberg Research Institute). The lab work was performed in the Grunelius-Mölgaard laboratory (Senckenberg Research Institute, Frankfurt am Main) with acknowledgments to Heike Kappes, Carola Grewe and Damian Baranski for lab support. We thank Annukka Korpijaakko, Natural Resources Institute Finland, for information about the DNA extraction of *Lecanora cadubriae* and *L. circumborealis* and acknowledge the DNA Sequencing and Genomics Laboratory, Institute of Biotechnology, University of Helsinki, for sequencing these samples. These two samples were sequenced with the support of Research Council of Finland, grant 1133858. A special thanks to Diego Morales-Briones for his assistance with the phylogenetic analyses and for engaging discussions. Felix Grewe and H. Thorsten Lumbsch kindly provided the genome sequence of *Lecanora helva* prior to publication. We are grateful to Rainer Cezanne, Marion Eichler and Einar Timdal as well as the curators of the New York Botanical Garden (NY), the Herbarium Mycologicum Academiae Sinicae-Lichenes, Beijing, China (HMAS-L), and the Lichen Herbarium of Kunming Institute of Botany, Chinese Academy of Sciences, Kunming, China (KUN-L) for providing a number of specimens included in this study.

## Author Contributions

Conceptualization, LL, CP, and JG; samples preparation and lab work, LL, CI, and EV; investigation, LL, CI, CP, and JG; methodology, software and formal analysis, JG; resources and funding acquisition, CP and LeM; project administration, CP; SL, LuM, LeM, SPO, and LW provided the samples; data curation and visualisation, LL and JG; writing—original draft preparation, LL, CP, and JG; writing—review and editing, LL, CI, CP, and JG; supervision, CP and JG. All authors contributed to the manuscript.

## Data availability

All the scripts, alignments and individual phylogenetic trees are available on figshare: <u>doi.XXX</u>. The raw data are deposited on Sequencing Read Archive (SRA) with the corresponding GenBank accession numbers listed in Table S1.

## Supplementary material

**File S1.** MultiQC report of the raw reads of the phylogenomic dataset used in this study.

**File S2.** Captus-assembly: Quality Control Report of the trimmed reads, which includes Summary Table, Stats of Reads/Bases, Per Base Quality, Per Read Quality, Read Length Distribution, Per Base Nucleotide Content, Per Read GC Content, Sequence Duplication Level, and Adapter Content.

**File S3.** Captus-assembly: Assembly Report of the *de novo* assembled metagenomes. The Summary Table shows the general assembly statistics for each sample. The Visual Stats plot shows more detailed statistics before and after filtering for each sample as a bar graph. The Length Distribution plot shows the distribution of contig lengths before and after filtering for each sample as a heatmap, and the Depth Distribution plot shows the distribution of contig depths before and after filtering for each sample as a heatmap.

**File S4.** Captus-assembly: Extract (Marker Recovery Report). The heatmap shows an extraction result for our *Lecanoraceae* dataset. The blue bars along with *x*- and *y*-axes indicate how many loci are recovered in each sample and how many samples each locus is recovered in, respectively.

**File S5.** SMRT Link User Guide v11.1. This document describes how to use PacBio’s SMRT Link software. SMRT Link is the web-based end-to-end workflow manager for Sequel II systems and Sequel II systems.

**File S6.** The coalescence-based phylogeny of the family *Lecanoraceae*, inferred from 4649 homologous genes using ASTRAL. Colours in the pie charts: blue denotes the proportion of concordant gene tree topologies, dark grey denotes the proportion of gene trees with the main alternative topology, red denotes the proportion of gene trees with the remaining discordant topologies, light grey denotes the proportion of gene trees with missing taxa and dark grey denotes the proportion of uninformative gene trees. The outgroup consists of *Gypsoplaca macrophylla*, *Letharia vulpina*, *Parmelia squarrosa*, and *Platismatia glauca*.

**Table S1.** Contents of the phylogenomic dataset used in this study. The table lists all taxa sampled in the phylogenomic analyses, together with specimen voucher and locality information, genome assembly identifiers or GenBank accession numbers, data sources, sequencing technologies, and an indication of generic type status where applicable. In total, 65 genome assemblies representing *Lecanoraceae* and selected outgroup taxa are included.

## REFERENCES

1. Anamoorthy S, Minh BQ, Wong TKF, et al. (2017). ModelFinder: fast model selection for accurate phylogenetic estimates. Nature Methods 14: 587–589. 10.1038/nmeth.4285

2. Andrews S (2010). FastQC: A quality control tool for high throughput sequence data. Available online: https://www.bioinformatics.babraham.ac.uk/projects/fastqc/

3. Arup U, Holien H, Coppins BJ (2023). *Lecanora caledonica* – a new species in the *Lecanora intumescens* group (*Lecanoraceae*) from north-western Europe. The Lichenologist 55: 107–114. 10.1017/S0024282923000233

4. Brodo IM, Haldeman M, Malíček J (2019). Notes on species of the *Lecanora albella* group (*Lecanoraceae*) from North America and Europe. The Bryologist 122: 430–450.

5. Chen YP, Su PW, Hyde KD, et al. (2023). Phylogenomics and diversification of *Sordariomycetes*. Mycosphere 14: 414–451. 10.5943/mycosphere/14/1/5

6. Choisy M (1929). Genres nouveaux pour la lichénologie dans le groupe des *Lécanoracées*. Bulletin de la Société Botanique de France 76: 521–527. 10.1080/00378941.1929.10837179

7. Davydov EA, Yakovchenko LS, Hollinger J, et al. (2021). The new genus *Pulvinora* (*Lecanoraceae*) for species of the ‘*Lecanora pringlei*’ group, including the new species *P. stereothallina*. The Bryologist 124: 242–256. 10.1639/0007-2745-124.2.24

8. Díaz-Escandón D, Tagirdzhanova G, Vanderpool D, et al. (2022). Genome-level analyses resolve an ancient lineage of symbiotic *ascomycetes*. Current Biology 32: 5209–5218. 10.1016/j.cub.2022.11.014

9. Eigler G (1969). Studien zur Gliederung der Flechtengattung *Lecanora*. Dissertationes Botanicae 4: 1–195.

10. Ekman S, Andersen HL, Wedin M (2008). The limitations of ancestral state reconstruction and the evolution of the ascus in the *Lecanorales* (lichenized *Ascomycota*). Systematic Biology 57: 141–156.

11. Elix JA, McCarthy PM (2017). Six new lichen species (*Ascomycota*) from Australia. Telopea 20: 147–163. 10.7751/telopea11598

12. Galindo LJ, López-García P, Torruella G, et al. (2021). Phylogenomics of a new fungal phylum reveals multiple waves of reductive evolution across Holomycota. Nature Communications 12: 4973. 10.1038/s41467-021-25308-w

13. González-Domínguez J, Schmidt B (2016). ParDRe: faster parallel duplicated reads removal tool for sequencing studies. Bioinformatics 32: 1562–1564. 10.1093/bioinformatics/btw038

14. Grewe F, Ametrano C, Widhelm TJ, et al. (2020). Using target enrichment sequencing to study the higher-level phylogeny of the largest lichen-forming fungi family: Parmeliaceae (Ascomycota). IMA Fungus 11: 27. 10.1186/s43008-020-00051-x

15. Grube M, Baloch E, Arup U (2004). A phylogenetic study of the *Lecanora rupicola* group (Lecanoraceae, Ascomycota). Mycological Research 108: 506–514. 10.1017/S0953756204009888

16. Hafellner J. 1984. Studien in Richtung einer natürlicheren Gliederung der Sammelfamilien *Lecanoraceae* und *Lecideaceae*. Nova Hedwigia, Beiheft 79: 241–371.

17. He MQ, Ca B, Liu F, et al. (2024). Phylogenomics, divergence times and notes of orders in Basidiomycota. Fungal Diversity 126: 127–406. 10.1007/s13225-024-00535-w

18. Hertel H (1983). Über einige aus *Lecidea* und *Melanolecia* auszuschließende Arten. Mitteilungen der Botanischen Staatssammlung München 19: 441–447.

19. Hertel H, Rambold G (1987). *Miriquidica* genus novum *Lecanoracearum* (*Ascomycetes* lichenisati). Mitteilungen der Botanischen Staatssammlung München 23: 377–392.

20. Hertel H, Rambold G. 1995. On the genus *Adelolecia* (lichenized Ascomycotina, Lecanorales). Bibliotheca Lichenologica 57: 211–230.

21. Hyde KD, Noorabadi MT, Thiyagaraja V, et al. (2024). The 2024 outline of fungi and fungus-like taxa. Mycosphere 15: 5146–6239.

22. Imshaug HA, Brodo IM (1966). Biosystematic studies on *Lecanora pallida* and some related lichens in the Americas. Nova Hedwigia 12: 1–59.

23. Ivanovich C, Weber L, Palice Z, et al. (2025a). A taxonomic revision of the lichen genus *Lecanoropsis* (*Lecanoraceae*). Phytotaxa 695: 1–56.

24. Ivanovich C, Weber L, Li L, et al. (2025b). New phylogenetic insights into the lichen genus Lecanora s.l. (Lecanoraceae, Ascomycota): resurrection of the genera Glaucomaria, Straminella and Zeora. The Lichenologist: In press.

25. Kalb K (1991). Lichenes Neotropici, Fascicle XII (Nos. 476–525). Neumarkt: 16 pp.

26. Kalb K (1994). *Frutidella*, eine neue Flechtengattung für *Lecidea caesioatra* Schaerer. Hoppea, Denkschrift der Regensburgischen Naturforschenden Gesellschaft 55: 581–586.

27. Kalyaanamoorthy S, Minh BQ, Wong TKF, et al. (2017). ModelFinder: fast model selection for accurate phylogenetic estimates. Nature Methods 14: 587–589.

28. Kandemir H, Dukik K, de Melo Teixeira M, et al. (2022). Phylogenetic and ecological reevaluation of the order *Onygenales*. Fungal Diversity 115: 1–71. 10.1007/s13225-022-00506-z

29. Kantvilas G (2016). Further additions to the lichen genus *Ramboldia* (*Lecanoraceae*) from Australia. Muelleria 34: 103–109. 10.5962/p.292271

30. Katoh K, Rozewicki J, Yamada KD (2019). MAFFT online service: multiple sequence alignment, interactive sequence choice and visualization. Briefings in Bioinformatics 20: 1160–1166. 10.1093/bib/bbx108

31. Kistenich S, Timdal E, Bendiksby M, et al. (2018). Molecular systematics and character evolution in the lichen family *Ramalinaceae* (*Ascomycota*: *Lecanorales*). Taxon 67: 871–904.

32. Kondratyuk SY, Lőkös L, Halda JP, et al. (2016). New and noteworthy lichen-forming and lichenicolous fungi 4. Acta Botanica Hungarica 58: 75–136.

33. Kondratyuk SY, Lőkós L, Jang S-H, et al. (2019). Phylogeny and taxonomy of *Polyozosia*, *Sedelnikovaea* and *Verseghya* of the *Lecanoraceae*. Acta Botanica Hungarica 61: 137–184. 10.1556/034.61.2019.1-2.9

34. Kozlov AM, Darriba D, Flouri T, et al. (2019). RAxML-NG: a fast, scalable and user-friendly tool for maximum likelihood phylogenetic inference. Bioinformatics 35: 4453–4455.

35. Lagou LJ, Kadereit G, Morales-Briones DF (2024). Phylogenomics uncovers rapid radiation and hybridization in *Cypripedium*. Annals of Botany 134: 1229–1250. 10.1093/aob/mcae161

36. Leuckert C, Poelt J (1989). Studien über die *Lecanora rupicola*-Gruppe in Europa (*Lecanoraceae*). Nova Hedwigia 49: 121–167.

37. Li Y, Steenwyk JL, Chang Y, et al. (2021). A genome-scale phylogeny of the kingdom Fungi. Current Biology 31: 1653–1665. 10.1016/j.cub.2021.01.074

38. Liimatainen K, Kim JT, Pokorny L, et al. (2022). A revised classification of *Cortinariaceae* based on genomic data. Fungal Diversity 112: 89–170. 10.1007/s13225-022-00499-9

39. Linder HP, Hardy CR, Rutschmann F (2005). Taxon sampling effects in molecular clock dating: an example from the African *Restionaceae*. Molecular Phylogenetics and Evolution 35: 569–582. 10.1016/j.ympev.2004.12.006

40. Lumbsch HT, Plümper M, Guderley R, et al. (1997). The corticolous species of *Lecanora* s.str. with pruinose apothecial discs. Symbolae Botanicae Upsalienses 32: 131–161.

41. Medeiros ID, Mazur E, Miadlikowska J, et al. (2021). Turnover of lecanoroid mycobionts and *Trebouxia* photobionts along an elevation gradient in Bolivia. Frontiers in Microbiology 12: 774839.

42. Miadlikowska J, Kauff F, Högnabba F, et al. (2014). A multigene phylogenetic synthesis for the class *Lecanoromycetes*. Molecular Phylogenetics and Evolution 79: 132–168.

43. Morales-Briones DF, Gehrke B, Huang C-H, et al. (2022). Analysis of paralogs in target enrichment data reveals ancient polyploidy in *Alchemilla* s.l. Systematic Biology 71: 190–207.

44. Ortiz EM, Höwener A, Shigita G, et al. (2023). A novel phylogenomics pipeline reveals reticulate evolution in *Cucurbitales*. bioRxiv. 10.1101/2023.10.27.564367

45. Palmer JM, Stajich J (2020). Funannotate: eukaryotic genome annotation. Zenodo. 10.5281/zenodo.4054262

46. Pease JB, Brown JW, Walker JF, et al. (2018). Quartet sampling distinguishes lack of support from conflicting support in the green plant tree of life. American Journal of Botany 105: 385–403.

47. Pérez-Ortega S, Turégano Y, Svensson M, et al. (2023). *Nimisora*, a new genus for a common lecideoid epiphyte from central Iberia. The Lichenologist 55: 335–345.

48. Poelt J (1983). *Bryonora*, eine neue Gattung der Lecanoraceae. Nova Hedwigia 38: 73–111.

49. Qu H, Cai Q, Redhead SA, et al. (2025). Suprageneric classifications in the genomic era: *Agaricineae*, *Pluteineae* and *Tricholomatineae*. Fungal Diversity 132: 127–150.

50. Rodriguez-Flakus P, Printzen C (2014). *Palicella*, a new genus of lichenized fungi and its phylogenetic position in the *Lecanoraceae*. The Lichenologist 46: 535–552. 10.1017/S0024282914000127

51. Schmull M, Miadlikowska J, Pelzer M, et al. (2011). Phylogenetic affiliations of members of the heterogeneous genus *Lecidea* s.lat. Mycologia 103: 983–1003.

52. Śliwa L, Mazur E, Wirth V (2023). Eine im phylogenetischen Kontext der Gattung bemerkenswerte neue *Myriolecis*-Art. Herzogia 36: 371–386.

53. Smith SA, Moore MJ, Brown JW, et al. (2015). Analysis of phylogenomic datasets reveals conflict, concordance and gene duplications. BMC Evolutionary Biology 15: 212.

54. Steenwyk JL, Buida TJ III, Li Y, et al. (2020). ClipKIT: a multiple sequence alignment trimming tool for phylogenomics. PLoS Biology 18: e3001007.

55. Svensson M, Westberg M (2021). A new lichenicolous species of *Carbonea* from Sweden. Phytotaxa 522: 221–230.

56. Svensson M, Haugan R, Timdal E, et al. (2022). The circumscription and phylogenetic position of *Bryonora*. Mycologia 114: 516–532.

57. Weber L, Ahti T, Ivanovich-Hichins C, et al. Contribution to the knowledge of lichens in Nyungwe National Park, Rwanda. Willdenowia: In review.

58. Yang Y, Smith SA (2014). Orthology inference in non-model organisms using transcriptomes and low-coverage genomes. Molecular Biology and Evolution 31: 2199–2215.

59. Zahlbruckner A (1926). Lichenes. In: Engler A (ed.), Die natürlichen Pflanzenfamilien, 2nd ed., Vol. 8: 61–270. Leipzig.

60. Zhang C, Scornavacca C, Molloy EK, et al. (2020). ASTRAL-Pro: species-tree inference despite paralogy. Molecular Biology and Evolution 37: 3292–3307.

61. Zhang Y, Clancy J, Jensen J, et al (2022). Providing scale to a known taxonomic unknown: a 70-fold increase in species diversity in a cosmopolitan nominal taxon. Journal of Fungi 8: 490.

62. Zhang Y, Yin Y, Wang L, et al. (2024). Two new species of *Rhizoplaca* (*Lecanoraceae*) from Southwest China. MycoKeys 101: 233–248.

63. Zhao X, Leavitt SD, Zhao ZT, et al. (2016). Towards a revised generic classification of lecanoroid lichens. Fungal Diversity 78: 293–304.

